# Cell segmentation without annotation by unsupervised domain adaptation based on cooperative self-learning

**DOI:** 10.1101/2024.07.05.602197

**Authors:** Shintaro Miyaki, Takashi Morikura, Shori Nishimoto, Yuta Tokuoka, Takahiro G Yamada, Akira Funahashi

## Abstract

Semantic cell segmentation from microscopic images is essential for the quantitative evaluation of cell morphology. Although supervised deep-learning-based models offer accurate segmentation, their performance degrades for unknown cell types. To address this problem, unsupervised domain adaptation methods based on adversarial training, self-training, or a combination of these approaches have been developed in recent years. These methods train the model using inference labels from the unknown domain as pseudo labels with reliability to resolve the discrepancy between the features of the unknown and known domains. However, conventional methods require a predefined threshold to calculate pseudo-labels reliability, leading to costly hyperparameter tuning. Here, we developed an unsupervised domain adaptation for semantic cell segmentation with cooperative self-learning (CULPICO: Cooperative Unsupervised Learning for PIxel-wise COloring) that does not require predefined threshold of the pseudo-labels reliability. The proposed method consists of two independent segmentation models and a mutual exchange mechanism of inference data. The models infer a label probability at each pixel and generate a pseudo-label as unsupervised learning. The pseudo-labels created by each model are mutually used as ground truth in the other model. Loss function is corrected by considering pixel-level discrepancies between the label probabilities inferred by the two models. The proposed method, despite being an unsupervised learning method, can segment efficiently the unknown cell types without labels with an accuracy comparable to supervised learning models. Our method, which could solve the performance degradation problem without constructing new datasets, is expected to accelerate life science by reducing the cost of extracting quantitative biological knowledge.

## Introduction

Recent advances in microscopy have made it possible to obtain large numbers of cellular microscopic images. For the acceleration of life science, it is essential to construct mathematical techniques to extract quantitative biological knowledge from such a huge amount of data [1]. In particular, semantic cell segmentation, which extracts cellular regions from microscopic images, is an important technique for quantifying morphological features of cells and gaining new insights into biological phenomena [2]. Traditional cell segmentation has been supported by image processing techniques such as image filtering and morphological operations [3]. However, those approaches require pre-determination of processing parameters for each image, resulting in a lack of objectivity and low throughput.

Recently, deep-learning-based algorithms that can automatically determine the parameters have been proposed [4, 5, 6, 7]. Deep-learning models that automate image processing operations while eliminating arbitrariness can solve the objectivity and throughput issues. Such models include convolutional neural networks (CNNs), which use convolutional processing, that are trained with microscopic images as input and a label matrix annotated with whether each pixel is a cellular region or not as output. These models generally consist of an encoder model that maps features from the input microscopic image to the latent space and a decoder model that outputs a label matrix from the latent representation. Previous studies have reported that supervised CNNs trained on certain cell types show better segmentation accuracy than the traditional image processing methods [4]. However, the segmentation performance of such supervised trained models is often degraded for unknown data not included in the training data [8]. A naive approach to this problem is to annotate the unknown cell types with additional correct labels. However, the cost of constructing a new dataset for each unknown cell type would be extremely high because the segmentation task requires pixel-level labeling in one image while general classification tasks require only image-level labeling. Learning methods that can achieve highly accurate segmentation of unknown cell types without additional annotation are required.

In deep learning techniques, it is generally assumed that feature distributions (domains) of training and inference data follow the same distribution. By contrast, if each domain were significantly different from the others, the inference performance would be degraded [8]. This phenomenon is usually called domain shift. Toward solving this problem, some algorithms based on unsupervised domain adaptation have been developed [9, 10, 11, 12]. There are currently two approaches in the field of unsupervised domain adaptation: adversarial learning and self-training.

In the adversarial learning approach, the models learn to align the feature distributions of the unknown target domain with those of the known source domain with the correct labels [13, 14, 15]. This approach tries to align the feature distributions by adversarial training based on the discrepancy between the feature distributions inferred by the models. One major issue with adversarial learning approaches is that, while aligning feature distributions between domains may be relatively easy, it is difficult to ensure the pixel-level consistency that is essential for improving segmentation performance. In addition, the adversarial learning algorithms themselves show unstable behavior during training [16, 17].

In the self-learning approach, the models learn to align the feature distribution by reusing the predicted labels for the target domain data as pseudo-labels [18, 19, 20]. Currently, mean teacher models, which utilize a student model that learns segmentation and a teacher model that generates pseudo-labels in the target domain, are widely used [18]. In general, the teacher model ‘s parameters are updated using an exponential moving average of the student model ‘s parameters. However, this approach introduces a coupling problem between the parameters of the teacher and student models, leading to performance degradation. To avoid this issue, Na et al. proposed a “dual teacher” model by constructing two teacher models for one student model and using them alternately [20]. However, one major challenge with these self-training approaches is that if the pseudo-labels used as correct labels are inaccurate, the model learning may be misdirected and will fail to improve performance. In addition, the approach of using multiple models to avoid the parameter coupling has the intrinsic problem that the number of parameters increases, which in turn increases the computational cost.

As another direction for improving the performance of unsupervised domain adaptation, an approach that integrates the adversarial training and self-training approaches has attracted attention. Wang et al. developed a new integrated approach referred to as “cross-region alignment” [21]. They first partitioned the target domain data into two regions, trusted and untrusted, based on whether the cross-entropy of the class probability at each pixel exceeded a pre-defined threshold (a hyperparameter). After the partitioning, they tried to align the feature distributions of these regions in the target domain by training the models so that they can accurately separate trusted and untrusted regions. This integrated approach utilizing the pixel-level reliability of pseudo-labels has been expected to improve the performance of unsupervised domain adaptation; however, as Wang et al. themselves pointed out, their approach is sensitive to the pre-defined threshold for calculating the discrete reliability [21].

Here, to improve the performance of unsupervised domain adaptation in the semantic cell segmentation task, we developed a cooperative self-learning framework using the pixel-level continuous reliability of pseudo-labels,referred to as discrepancy. Our architecture consists of two independent encoder–decoder models. In our approach, while both models learn the source domain data, the labels predicted by each model in the target domain data are used as pseudo-labels for each other ‘s training. To avoid instability in training caused by errors in the pseudo-labels, the pixel-level differences in the probabilities of the predicted labels are used during the self-training using pseudo-labels. The main contribution of this study is that we introduce a concept of continuous pixel-level reliability (discrepancy) that does not require prior definition and present a new integrated unsupervised domain adaptation approach in which each model simultaneously plays the role of both the teacher model and the student model, called cooperative self-learning. To assess the performance of our method, the segmentation accuracy was evaluated by a public dataset of cell images acquired with phase-contrast microscopy. The proposed method robustly improved the segmentation performance even for unknown cell types. Notably, the proposed method, despite being an unsupervised learning method, outperformed the performance of a supervised learning model for some cell types. A model analysis indicated that the simultaneous updating of pseudo-labels and models considering the pixel-level discrepancies may have contributed to improving the performance by suppressing false negatives. The proposed method, which achieves segmentation accuracy as high as supervised learning even for unknown cell types without correct labels, is expected to accelerate the development of life science by reducing the cost of extracting quantitative biological knowledge.

## Methods

### Dataset

To assess the performance of our method, we used a public dataset of cell images published by Edlund et al. [22]. The dataset consists of phase-contrast microscopic images of eight cell types showing a variety of cell morphologies, and label images annotated with cellular and background regions by experts. In this study, the models were trained with the phase-contrast microscopic images as input and the labeled images as output. Representative microscopic images of the eight cell types are shown in Supplementary Fig.S1. The cell types are human glioblastoma cell line A172, human breast cancer cell line BT-474, mouse macroglial cell line BV-2, human hepatocellular carcinoma cell line Huh7, human breast cancer cell line MCF7, human neuroblastoma cell line SH-SY5Y, human breast cancer cell line SK-BR-3, and human ovarian cancer cell line SK-OV-3. Detailed information on image acquisition can be found in the paper by Edlund et al. [22].

Because the purpose of this study is to develop a cell morphology segmentation algorithm, data with missing cell region labels or data with extremely high cell density were excluded from the dataset in this study (Supplementary Fig.S2). In addition, to minimize the effects of well plate–derived characteristic image patterns on learning, we constructed training and test datasets using microscopic images of cells cultured in different well plates, respectively (Supplementary Table S1). The training data were split 3:1 into training data and validation data.

To analyze morphological features of each cell type, microscopic images were cropped to 224 × 224 pixels from the image center. After the cropping, feature representations with 4096 dimensions were extracted using the VGG16 model [23], which was trained for the image classification task on ImageNet [24], a benchmark dataset for natural images. To qualitatively evaluate the morphological feature distribution, representations were reduced to two dimensions using t-SNE [25] and then plotted in two-dimensional space.

### Overview of the proposed method

The proposed method (CULPICO: Cooperative Unsupervised Learning for PIxel-wise COloring) consisted of two independent encoder–decoder models with a mutual exchange mechanism (Fig. 1). To achieve cooperative self-learning, we designed a loss function using the pixel-level discrepancy. The encoder–decoder model was constructed by a U-Net architecture consisting of different initial value parameters (Supplementary Fig. S3). In the U-Net architecture, batch normalization was performed after each convolution or inverse convolution process.

**Figure 1:**
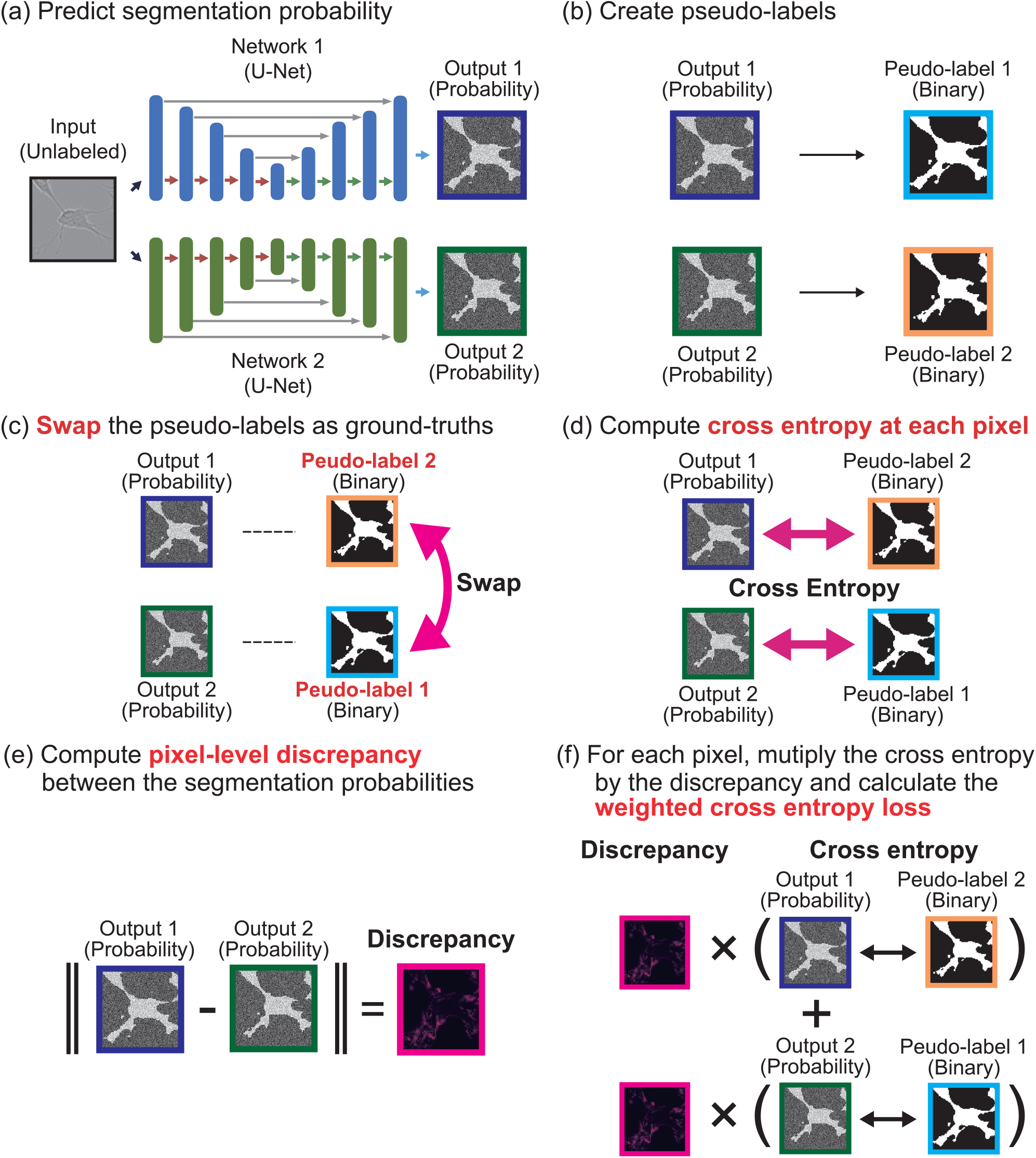
Schematic diagram of the proposed method for performing collaborative self-learning on the target dataset. (a) Infer label probabilities for the input images on Network 1 and Network 2, respectively. (b) Generate pseudo-labels from the inferred label probabilities by binarizing based on a pre-set threshold value. (c) The pseudo-label generated in one model is used as the pseudo-correct label in the other model. (d) Compute the cross entropy between the inferred label probability and the pseudo-correct label for each pixel. (e) Compute the pixel-level discrepancy using the absolute difference between the inferred label probabilities at each pixel. (f) Compute a weighted cross entropy multiplied by the discrepancy for each pixel. The specific formulas of the loss function are explained in the Methods section.

In this study, we defined the datasets consisting of microscopic images with labels as the source dataset and the datasets consisting of microscopic images without labels as the target dataset. The proposed method achieves unsupervised domain adaptation by simultaneously performing supervised learning on the source dataset and cooperative self-learning on the target dataset.

When training on the source domain data, supervised learning was performed using cross-entropy as the loss function. For the target domain data, each U-Net model predicted a label probability map for the unlabeled input images (Fig. 1a). Pseudo-labels were then generated by binarizing the label probability map using a predefined threshold (Fig. 1b). The generated pseudo-labels were mutually used as pseudo ground truth labels for each U-Net model (Fig. 1c). Next, the cross-entropy between the pseudo-labels and the predicted label probability maps was calculated on a per-pixel basis (Fig. 1d). Since the pseudo-labels contain uncertainty in the inference, the absolute differences between the predicted label probability maps were computed as a discrepancy for each pixel (Fig. 1e). By multiplying the cross-entropy and the discrepancy at the pixel level, a weighted cross-entropy was obtained (Fig. 1f). Finally, the total error was calculated by summing the cross-entropy computed from the source domain data and the weighted cross-entropy computed from the target domain data. The parameters of each U-Net model were updated by back-propagating the total error, using the Adam algorithm with a learning rate of 0.001.

For the supervised learning on the source dataset, we trained the models using the microscopic image as input and the labels as output. The loss function *L*_*s*_ of the supervised learning is defined as follows based on the cross-entropy error between the inferred label probability and the correct label.

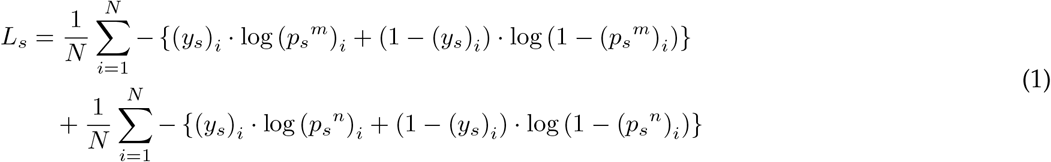

where *i* is the index number of each pixel and *N* is the total number of pixels in the image. The (*p*_*s*_)_*i*_ is *i*-th pixel in the inferred label probability for the image and (*y*_*s*_)_*i*_ is *i*-th pixel in the correct label. The *m* and *n* denote different encoder-decoder models. For the cooperative self-learning on the target dataset, the models inferred label probabilities representing the probability of cellular regions in the target image, and pseudo-labels were generated by binarizing them with a predefined threshold value. The pseudo-label generated in one model was then used as the pseudo-correct label in the other model. The loss function *L*_*t*_ of the cooperative self-learning was designed based on the weighted cross-entropy error according to pixel-level discrepancy between the inference label probabilities in the two models (Eq. (2)). The loss function *L*_*t*_ was expected to actively learn the error in pixels with low discrepancy, while carefully learning the error in pixels with high discrepancy.

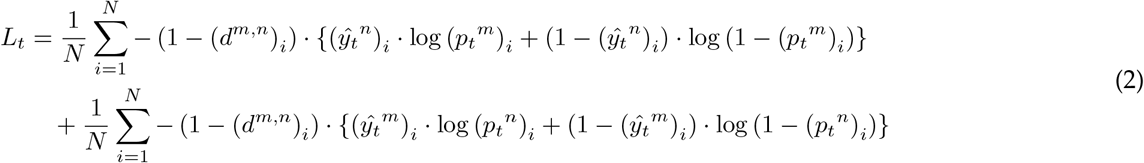

where (*p*_*t*_)_*i*_ is *i*-th pixel in the label probability inferred for the target image, the (*ŷ*_*t*_)_*i*_ is the *i*-th pixel in the pseudo-correct label (Eq. (3)), and the (*d*^*m*,*n*^)_*i*_ is the pixel-level discrepancy between the label probabilities inferred from the two models *m* and *n* (Eq. (4)).

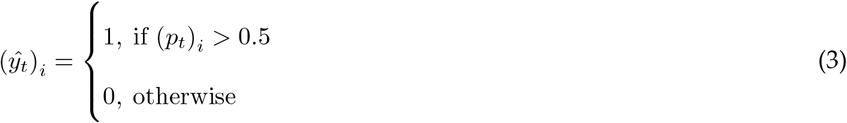

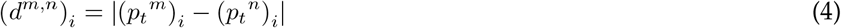

where label 1 is the cellular region and label 0 is the background region.

Using the above loss functions *L*_*s*_ and *L*_*t*_, the total loss function *L*_*total*_ was defined as follows (Eq. (5))

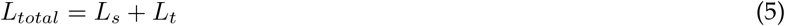

By performing the cooperative self-learning in the target dataset simultaneously with supervised learning in the source dataset, the feature distribution of the target dataset is expected to adapt to that of the source. Although previous methods were developed to improve the performance of segmentation by using errors at the image level [26], the proposed method was designed to use the errors at the pixel level.

### Evaluation metrics of segmentation

To evaluate the effectiveness of the proposed method, we developed a U-Net model as a lower-bound model that was trained using only the source dataset. In addition, we developed a U-Net model as an upper-bound model that was trained using only the target dataset. The lower-bound model is expected to degrade inference performance at the target because the target cell types are not included in the training data. The upper-bound model is equivalent to a general supervised learning model, and the inference performance at the target is expected to be relatively high because the target cell types are included in the training data. To compare our model to previous unsupervised domain adaptation models, we also trained Saito et al. ‘s model [15], which is an adversarial learning method; Zou et al. ‘s model [19], which is a self-learning method; Na et al.’s model [20], which is a recent mean teacher model; and Wang et al. ‘s model [21], which is an integrated adversarial learning and self-learning method. Saito et al. developed an architecture with two decoder models branching from a single encoder model [15]. This approach tries to align the feature distributions by adversarial training based on the image-level discrepancy between the feature distributions of the images. Zou et al. developed a single encoder–decoder model to generate pseudo-labels for unknown data without labels [19]. They tried to implicitly align the feature distributions by reusing the pseudo-labels as correct labels. Na et al. developed a dual teacher–student model that utilizes one student model that learns segmentation and two teacher models that generate pseudo-labels in the target domain [20]. In general teacher–student models, the weights of the teacher models are updated based on the exponential moving average of the weights of the student model [18]. Unlike the general approaches, they tried to align the feature distributions of the source and target domain by using two teacher models alternately to avoid coupling the weights of the teacher and student models. Wang et al. developed a new integrated approach referred to as “cross-region alignment” [21]. In cross-region alignment, the predicted images in the target domain were partitioned into trusted and untrusted regions. The partitioning was performed by the discrete reliability that was defined with a pre-defined threshold. If the cross-entropy of the class probability at each pixel exceeded the threshold, the pixel was included in the trusted regions. After the partitioning, they tried to align the feature distributions of these regions in the target domain by training a generator model so that it can accurately separate the trusted and untrusted regions, and to train the discriminator model so that it can accurately identify the trusted and untrusted regions. In the architecture of the previous models, batch normalization was performed after each convolution or inverse convolution process.

The training was performed using a GPU (NVIDIA V100 or A100) with a mini-batch size of 16 for 400 epochs. The microscopic images were normalized to [0,1] by performing min–max normalization. In addition, the microscopic images in which cells were located at the periphery of the image so that their entire cellular shape could not be captured were interpolated by mirror padding. The data were augmented by inverting left and right, ?ipping up and down, and random cropping. The image size after cropping was fixed at 352 × 272 pixels. The model at the epoch with the smallest loss to the data for validation was selected as the best model. We evaluated the performance of the trained model with the test data that were not used in training.

As an evaluation metric, we calculated the intersection-over-union metric (IoU), which is widely used to evaluate the performance of segmentation (Eq. (6)).

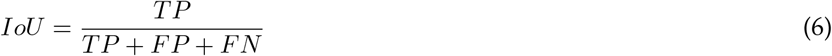

where *TP* , *FP* , and *FN* are the number of pixels counted as True Positive, False Positive, and False Negative, respectively. The range of IoU is from 0 to 1, and a value closer to 1 indicates better segmentation. In addition, we also calculate the Dice coefficient (DIC), which is another popular metric used in image segmentation (Eq. (7)).

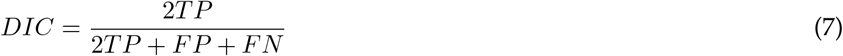

Dice coefficient is equivalent to F-measure metric, the harmonic mean between sensitivity and precision. DIC shows the balance between false positives and false negatives. The range of DIC is from 0 to 1, and a value closer to 1 indicates better segmentation. Although IoU and DIC are similar, IoU penalizes under- and over-segmentation more than DIC [27]. To evaluate the similarity of object boundary shape, we also calculated the 95% Hausdorff distance (HD95) and average surface distance (ASD). HD95 measures the 95th percentile of distance between the surface points of the ground truth object and the predicted object. ASD measures the average distance between the surface points of the ground truth object and the predicted object. HD95 evaluates the error of the worst 95% of the boundary shape, whereas ASD evaluates the average error of the boundary shape.

To calculate performance efficiency, we compared the average of segmentation performance and the number of parameters each model consisted of. A model with fewer parameters and higher performance is considered more efficient. To quantify the performance efficiency, we calculated each model ‘s average performance with IoU, DIC, HD95, and ASD for all combinations of cell types. After the calculation of the average, we applied z-score normalization to each of the average performance metrics and the number of parameters for each model. Then, we calculated the Euclidean distance from the upper-bound model for the normalized performance and parameter count. After the calculation of the Euclidean distance, we performed min–max normalization for the Euclidean distance. The resulting normalized Euclidean distance indicates the similarity relative to the upper-bound model, with values in the range [0, 1] where a value closer to 0 denotes more similarity to the upper-bound model. Finally, by subtracting the similarity from 1, we obtained a quantitative measure of performance efficiency. This performance efficiency metric, which also lies in the range [0, 1], indicates better efficiency as it approaches 1, signifying more similarity to the upper-bound model.

## Results

### Proposed method robustly improved the segmentation performance for unknown cell types

To establish an unsupervised domain adaptation method that can achieve highly accurate segmentation of unknown cell types without labels, we developed a cooperative self-learning method using the pixel-level discrepancy between different model inferences (Fig. 1). To evaluate the proposed method, we used the public dataset of cell images published by Edlund et al. [22]. The dataset consists of phase-contrast microscopic images of eight different cell types. In this study, we defined the source as the cell type for which the correct label is annotated, and the target as an unknown cell type for which the correct label is not annotated. We evaluated the performance of the proposed model using 56 different combinations of cell types used as source and target. To evaluate the effectiveness of the proposed method, we used as the lower-bound model a supervised U-Net model in which only the source cell type was used for training, and as the upper-bound model one in which the target cell type was used for training. In this study, the performance of the lower-bound model is the minimum one that must be achieved, and that of the upper-bound model is the target one.

To analyze the morphological characteristics of each cell type, the result of dimensionality reduction of the microscopic images is shown in Supplementary Fig. S4. In the reduced 2D space, it was shown that clusters were formed for each cell type. The upper-bound model, which infers the same cell types as in training, has the potential to achieve highly accurate segmentation, while the lower-bound model, which infers cell types different from those in training, may degrade the model performance due to the domain shift.

To evaluate the effect of cell-type differences during training and inference on the performance of the models, we trained 56 supervised U-Net models with different combinations of cell types. The performance of each U-Net model is shown in Table 1. The results showed that the upper-bound model has high accuracy for most of the combinations except for SH-SY5Y cells, but the lower-bound model showed lower accuracy than the upper bound model. The combinations with Huh7 cells as the target showed the most significant performance degradation among all cell types. Notably, the IoU of the lower-bound model with Huh7 cells as the target and BV-2 cells as the source was reduced by a maximum of 0.819 (0.867 → 0.048) relative to the upper bound model. The lower-bound model also showed that there were variations in IoU for each source cell type even if the target cell type was the same.

**Table 1:** Segmentation performance for each cell type using each U-Net model The vertical series represents the cell types used to train the model (source), and the horizontal series represents the cell types used to test the model (target). The values in the table show the mean IoU for the test data, and those in parentheses show the standard deviation. The IoUs of the upper bound models are underlined.

| Source \ Target | A172 | BT-474 | BV-2 | Huh7 | MCF7 | SH-SY5Y | SK-BR-3 | SK-OV-3 |
| --- | --- | --- | --- | --- | --- | --- | --- | --- |
| A172 | <u>0.878</u><br>(0.029) | 0.790<br>(0.078) | 0.804<br>(0.083) | 0.664<br>(0.063) | 0.858<br>(0.067) | 0.721<br>(0.066) | 0.865<br>(0.024) | 0.486<br>(0.104) |
| BT-474 | 0.910<br>(0.030) | <u>0.817</u><br>(0.055) | 0.827<br>(0.072) | 0.795<br>(0.057) | 0.810<br>(0.089) | 0.759<br>(0.064) | 0.887<br>(0.036) | 0.847<br>(0.048) |
| BV-2 | 0.161<br>(0.153) | 0.212<br>(0.237) | <u>0.842</u><br>(0.064) | 0.048<br>(0.044) | 0.291<br>(0.237) | 0.410<br>(0.112) | 0.745<br>(0.063) | 0.093<br>(0.054) |
| Huh7 | 0.915<br>(0.027) | 0.771<br>(0.099) | 0.795<br>(0.110) | <u>0.867</u><br>(0.054) | 0.731<br>(0.192) | 0.738<br>(0.054) | 0.889<br>(0.029) | 0.825<br>(0.084) |
| MCF7 | 0.902<br>(0.027) | 0.777<br>(0.072) | 0.813<br>(0.077) | 0.711<br>(0.060) | <u>0.847</u><br>(0.063) | 0.739<br>(0.062) | 0.866<br>(0.033) | 0.660<br>(0.098) |
| SH-SY5Y | 0.493<br>(0.095) | 0.375<br>(0.184) | 0.838<br>(0.061) | 0.156<br>(0.571) | 0.571<br>(0.167) | <u>0.789</u><br>(0.052) | 0.789<br>(0.053) | 0.575<br>(0.085) |
| SK-BR-3 | 0.918<br>(0.026) | 0.826<br>(0.080) | 0.834<br>(0.076) | 0.864<br>(0.055) | 0.863<br>(0.070) | 0.693<br>(0.085) | 0.903<br>(0.035) | 0.895<br>(0.037) |
| SK-OV-3 | 0.917<br>(0.034) | 0.785<br>(0.098) | 0.787<br>(0.089) | 0.837<br>(0.070) | 0.858<br>(0.074) | 0.765<br>(0.071) | 0.876<br>(0.051) | <u>0.917</u><br>(0.030) |

To evaluate the effectiveness of the proposed method, the proposed models were trained with 56 different combinations of source and target cell types. Representative learning curves of the proposed method are shown in Supplementary Fig. S5. At the end of the learning process, the error differences between the two models converged as the training progressed, suggesting that the two models were able to learn cooperatively. The results of the qualitative evaluation of the proposed model performance are shown in Fig. 2, and the results of the quantitative evaluation are shown in Table 2. Qualitatively, while the lower bound models frequently produced false negatives, the proposed model overcame the false-negative errors in the target domain. However, the proposed model also tended to produce false positives. These qualitative evaluations suggest that the proposed method was inclined to mitigate performance degradation due to domain shift by actively detecting cellular regions. In the quantitative evaluation, the proposed models had higher accuracy than the lower-bound model in 29 combinations (Table 2). It should be noted that the proposed models improved the IoU by 0.652 (0.156 → 0.808) compared with the lower-bound model in the combinations with Huh7 cells as the target, which showed the largest performance degradation between the upper-bound and lower-bound models, and SH-SY5Y cells as source, which showed the lowest accuracy in the upper-bound model. In addition, the proposed method showed higher accuracy than the lower-bound model when BT-474, Huh7, or SK-OV3 cells were used as the target cells. Furthermore, the proposed method showed higher accuracy than the lower-bound model when BV-2 or SH-SY5Y cells were used as the source cells. In contrast, the proposed method tended to show lower accuracy than the lower-bound model, especially when SH-SY5Y and BV-2 cells were used as the target. These results suggested that SH-SY5Y and BV-2 cells contain necessary shape information to construct cell shape features, while BT-474, Huh7, and SK-OV3 cells contain sufficient shape information to construct cell shape features. Taken together, these results indicate that the proposed method robustly improves the segmentation performance for unknown cell types compared with the lower-bound model.

**Table 2:** Segmentation performance for each cell type using the proposed method The vertical series represents the cell types used to train the model (source), and the horizontal series represents the cell types used to test the model (target). Values in the table show the mean IoUs for the test data, and those in parentheses show the standard deviation. Those in bold type represent the models with higher IoU than the lower-bound model.

| Source \ Target | A172 | BT-474 | BV-2 | Huh7 | MCF7 | SH-SY5Y | SK-BR-3 | SK-OV-3 |
| --- | --- | --- | --- | --- | --- | --- | --- | --- |
| A172 | - | 0.784<br>(0.074) | 0.518<br>(0.079) | **0.792**<br>(0.069) | 0.857<br>(0.068) | 0.617<br>(0.082) | 0.837<br>(0.052) | **0.884**<br>(0.051) |
| BT-474 | 0.852<br>(0.038) | - | 0.779<br>(0.068) | **0.861**<br>(0.058) | **0.861**<br>(0.067) | 0.712<br>(0.078) | **0.891**<br>(0.031) | **0.875**<br>(0.054) |
| BV-2 | **0.860**<br>(0.036) | **0.814**<br>(0.079) | - | **0.579**<br>(0.055) | **0.854**<br>(0.069) | **0.692**<br>(0.071) | **0.875**<br>(0.031) | **0.247**<br>(0.060) |
| Huh7 | 0.909<br>(0.030) | **0.820**<br>(0.078) | 0.686<br>(0.142) | - | **0.856**<br>(0.072) | 0.672<br>(0.074) | 0.879<br>(0.041) | **0.912**<br>(0.031) |
| MCF7 | 0.820<br>(0.045) | **0.819**<br>(0.080) | 0.786<br>(0.068) | **0.806**<br>(0.060) | - | 0.630<br>(0.080) | **0.872**<br>(0.029) | 0.632<br>(0.071) |
| SH-SY5Y | **0.907**<br>(0.028) | **0.814**<br>(0.078) | 0.827<br>(0.065) | **0.808**<br>(0.061) | **0.863**<br>(0.070) | - | **0.881**<br>(0.029) | **0.813**<br>(0.060) |
| SK-BR-3 | 0.912<br>(0.028) | 0.824<br>(0.079) | 0.826<br>(0.088) | 0.850<br>(0.059) | 0.859<br>(0.071) | **0.719**<br>(0.079) | - | 0.569<br>(0.064) |
| SK-OV-3 | 0.911<br>(0.037) | **0.806**<br>(0.080) | **0.800**<br>(0.092) | **0.844**<br>(0.061) | 0.854<br>(0.071) | 0.710<br>(0.075) | 0.868<br>(0.048) | - |

**Figure 2:**
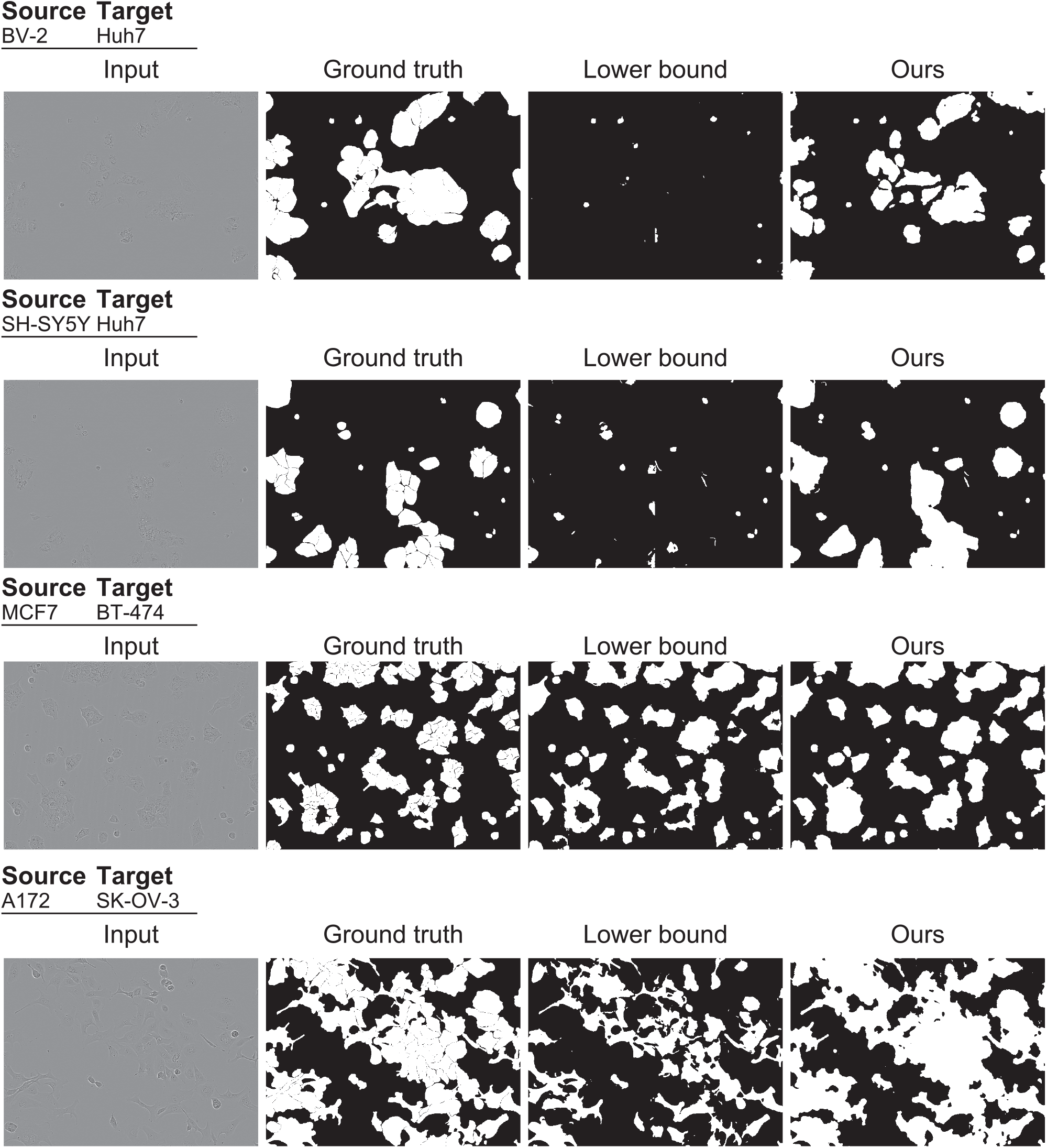
Representative segmentation results of the lower-bound model and the proposed model (ours) The columns show the input images, the ground truth images, the labels predicted by the lower bound model, and the labels predicted by the proposed method, respectively. The rows show the results when the source and target cell type combinations are BV2 → Huh7, SH-SY5Y → Huh7, MCF7 → BT-474 and A172 → SH-SK-OV-3, respectively.

### Proposed method, despite being an unsupervised learning model, outperforms segmentation of some unknown cell types compared with supervised learning

To evaluate the effectiveness of the proposed method, we compared and assessed the model performance of lower-bound models, upper-bound models, the adversarial learning model by Saito et al. [15], the self-learning model by Zou et al. [19], he dual teacher model by Na et al. [20], and the integrated adversarial learning and self-learning model by Wang et al. [21]. As evaluation metrics for segmentation performance, we used the IoU, DIC, HD95, and ASD. IoU and DIC evaluate the accuracy of object shape and object detection, whereas HD95 and ASD evaluate the accuracy of the object ‘s outline shape. While the DIC represents the precision of object detection by evaluating the balance between false negatives and false positives in segmentation, IoU represents the precision of object shape more strictly than the DIC. HD95 represents the local similarity between each point on the contour of objects, and ASD represents the global similarity of the contour of objects.

Qualitative evaluation of the performance comparisons is presented in Fig. 3, and the quantitative evaluation results are shown in Supplementary Tables S2-S5. In Saito et al. ‘s adversarial learning model, false-positive artifacts resembling high-frequency noise were observed. In contrast, in Zou et al. ‘s self-learning model, many false-negative errors were observed, similar to those observed in the lower-bound model. The model proposed by Na et al., which enhanced the self-learning approach, achieved the highest segmentation performance among the conventional models; however, noise-like false negatives were observed within the regions predicted as cellular regions. Wang et al. ‘s model, which integrated both adversarial and self-learning approaches, was expected to combine the strengths of each; nevertheless, its predictions exhibited both false-positive artifacts and noticeable false-negative errors. Conversely, the proposed method, similar to the upper-bound model, suppressed the false-positive artifacts and reduced the false-negative errors compared to the conventional models. Moreover, the noise-like false negatives within cellular regions that were observed in Na et al. ‘s model were also mitigated by the proposed method. In the quantitative evaluation, Na et al. ‘s model showed the highest performance among the conventional models and across the widest range of cell type combinations for IoU and DIC (IoU: 23/56, DIC: 24/56), which indicate performance in terms of object detection and shape, with the proposed model ranking second (IoU: 15/56, DIC: 15/56) (Supplementary Table S2, S3). In addition, Na et al. ‘s model exceeded the performance of the upper-bound model in the widest combination of cell types (IoU: 15/56, DIC: 15/56), with the proposed model ranking second (IoU: 14/56, DIC: 15/56). In terms of performance related to object contours, the lower-bound model showed the highest performance for the widest range of cell type combinations (HD95: 25/56, ASD: 23/56), followed by Saito et al. ‘s model (HD95: 11/56) and Zou et al. ‘s model (ASD: 12/56) (Supplementary Table S4, S5). The number of combinations of cell types that exceeded the upper-bound models of HD95 and ASD was the highest for the model of Saito et al. (HD95: 14/56, ASD: 11/56), followed by the model by Zou et al. (HD95: 12/56, ASD: 9/56). Considering the characteristics of these evaluation metrics, Na et al. ‘s model and the proposed model can accurately predict object detection and shape even in the target domain, while the lower-bound model, Saito et al. ‘s model, and Zou et al. ‘s model can predict contour shapes similar to the ground truth. Most importantly, even though the proposed method is an unsupervised learning model, it not only demonstrated higher accuracy than some conventional models, but also achieved accuracy comparable to the upper-bound model for certain combinations of cell types (combinations with A172, BT-474 and MCF7 cells as the target) (Supplementary Table S2). It was noteworthy that the proposed method showed higher accuracy than the upper-bound model when the source and target cell type combinations were BV-2 → MCF7, while the previous models showed lower accuracy than the upper-bound model. In contrast, when SH-SY5Y cells were used as the target, the proposed method showed lower accuracy than the upper-bound model, as in the previous models. Although the proposed method learned sufficient features for cell shape in A172, BT-474 and MCF7 cells, like the previous models it had difficulty learning the necessary features for cell shape in SH-SY5Y cells.

**Figure 3:**
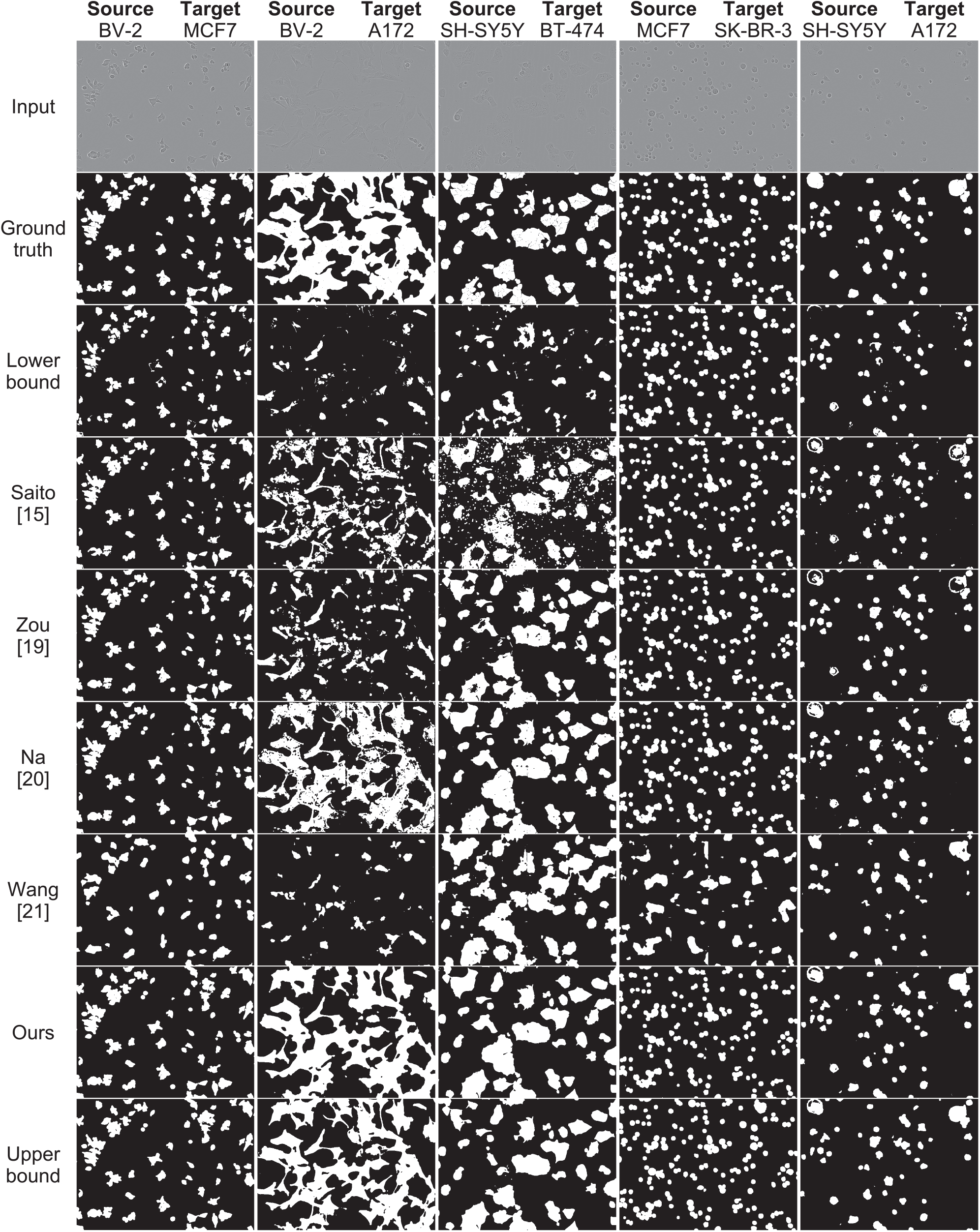
Representative segmentation results of lower-bound models, upper-bound models, conventional models, and ours The columns show the results when the source and target cell type combinations are BV2 → MCF7, BV2 → A172, SH-SY5Y → BT-474, MCF7 → SK-BR-3 and SH-SY5Y → A172, respectively. The rows show the input images, the ground truth images, and the labels predicted by the lower-bound model, the models of Saito[15], Zou [19], Na [20], Wang [21], the proposed method, and the upper-bound model, respectively.

Finally, we compared the model performances and the number of parameters, and evaluated the performance efficiency quantitatively (Fig. 4). With regard to IoU and DIC, although Na et al. ‘s model achieved the highest accuracy in object detection (Supplementary Table S2, S3), the proposed model demonstrated the best performance efficiency (Fig. 4a, Fig. 4b). While Na et al. ‘s model requires the construction of three models (one student model and two teacher models), the proposed model only requires the construction of two models, enabling it to achieve highly efficient segmentation. In terms of the performance efficiency for HD95 and ASD, the proposed model performed comparably to conventional models (Fig. 4c, Fig. 4d). Taken together, these results indicate that the proposed model cannot only achieve high-precision unsupervised segmentation with high efficiency, but can also achieve accuracy comparable to that of the upper-bound model in certain combinations of cell types.

**Figure 4:**
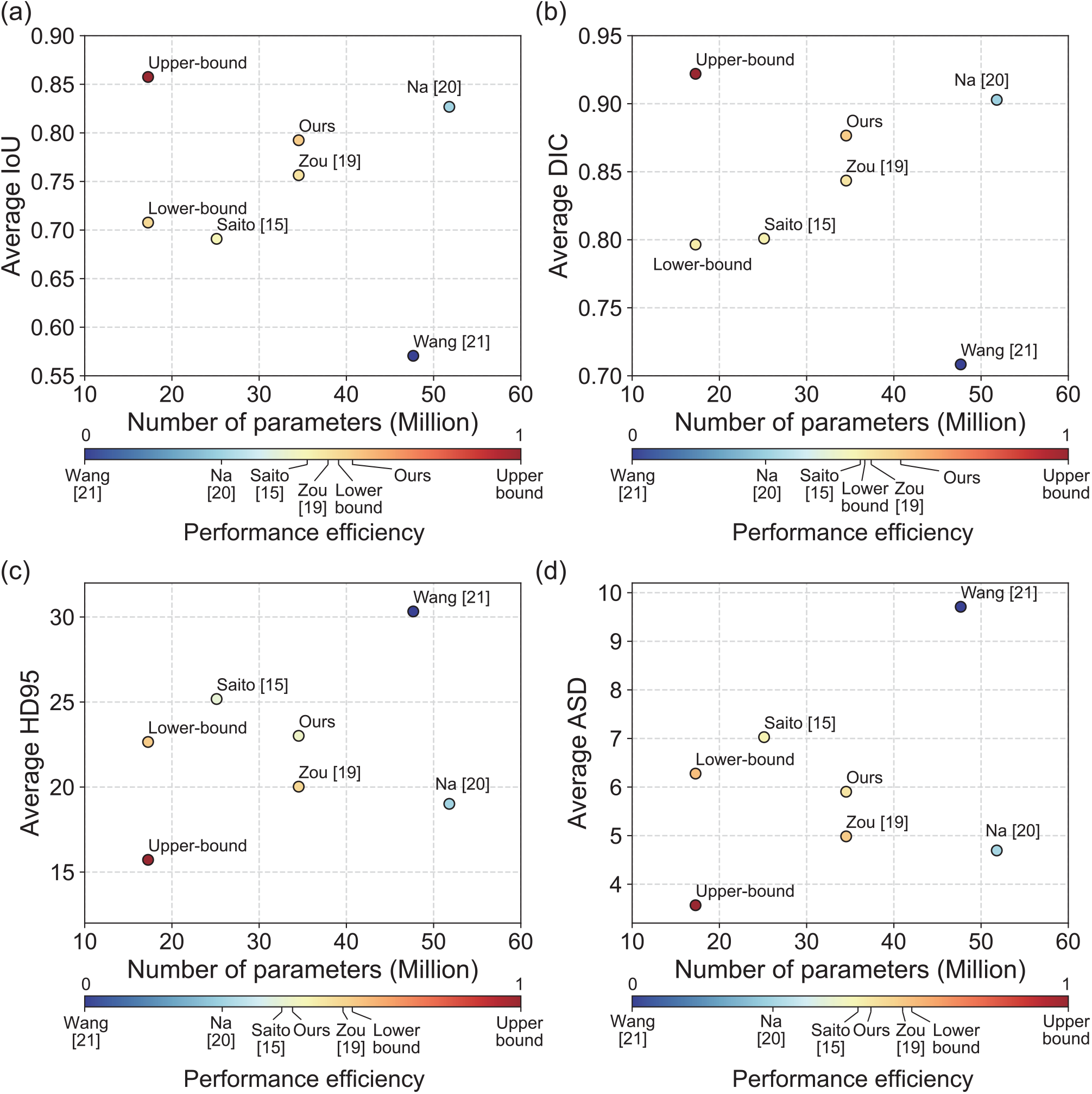
Comparison of the segmentation performance and number of model parameters The vertical axes represent the average segmentation performance of each metric, and the horizontal axes represent the number of model parameters. The plotted points represent each model. The text placed near each point represents each model ‘s name. The color bars represent the performance efficiency as defined in the Methods section. The range of performance efficiency is from 0 to 1, and a value closer to 1 indicates better performance efficiency. (a) IoU: The model plotted in the upper left is the most efficient model. (b) DIC: The model plotted in the upper left is the most efficient model. (c) HD95: The model plotted in the lower left is the most efficient model. (d) ASD: The model plotted in the lower left is the most efficient model.

### Simultaneous updating of the pseudo-labels and model to consider the pixel-level discrepancy might have improved the accuracy of detecting cellular regions

To verify the factors that contributed to the performance improvement of the proposed method, we qualitatively evaluated the segmentation results and verified the transition of the pixel-level discrepancy and pseudo-label accuracy during the training. Representative segmentation results are shown in Fig. 5. In the combination of SH-SY5Y → MCF7 (lower-bound: IoU 0.571 ± 0.167, Ours: IoU 0.863 ± 0.070) and SH-SY5Y → Huh7 (lower-bound: IoU 0.156 ± 0.571, Ours: IoU 0.808 ± 0.061), the true-positive regions increased and the false-negative regions decreased, indicating the performance improvement. In contrast, the combination of Huh7 → SH-SY5Y (lower-bound: IoU 0.738 ± 0.054, Ours: IoU 0.672 ± 0.074) did not suppress false-negative errors in the regions where protrusion-like structures were present. Taken together, these results suggested that the Huh7 and MCF7 cells contained sufficient features to construct cell shape, whereas the SH-SY5Y cells contained the necessary features to construct cell shape.

**Figure 5:**
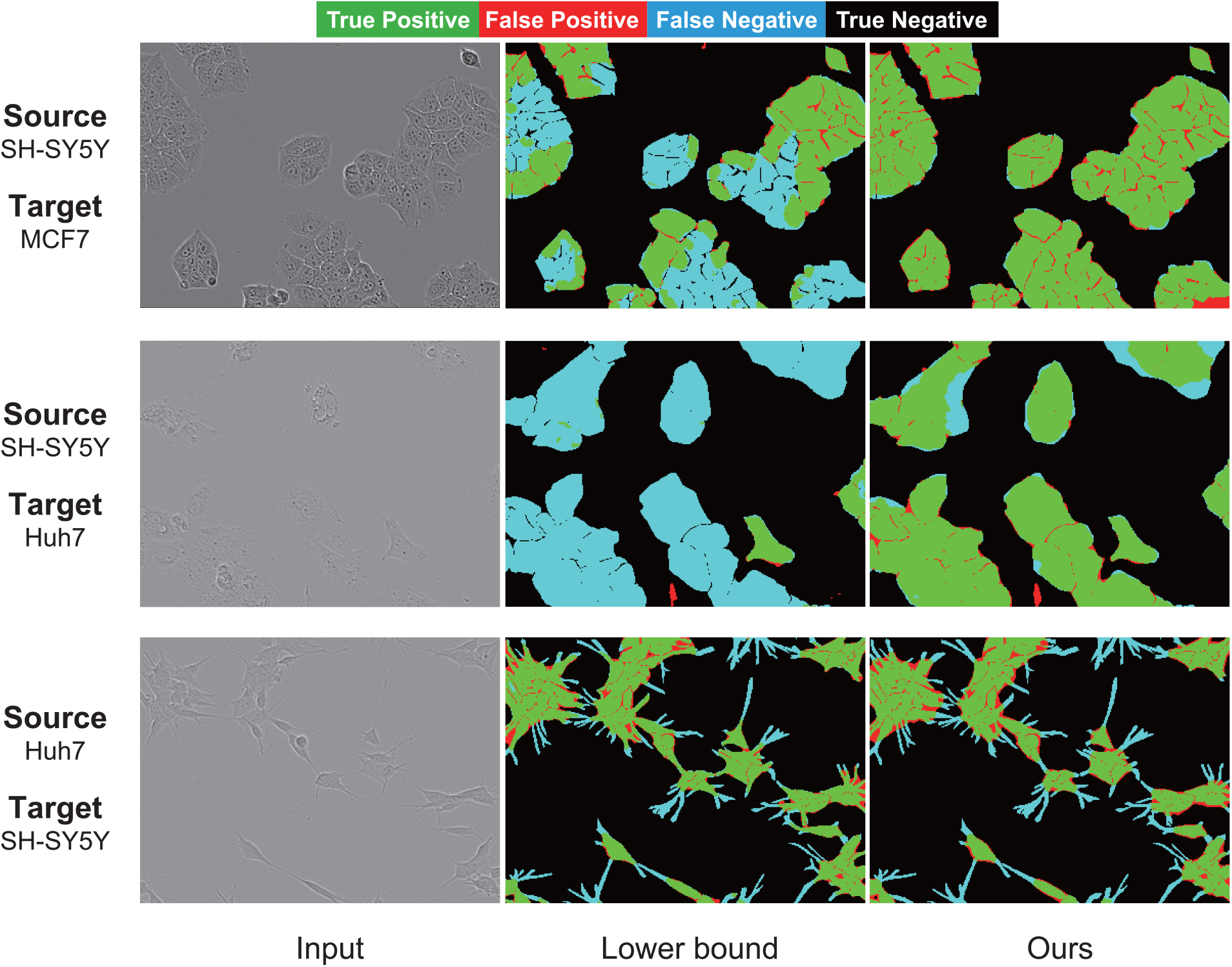
Representative errors in segmentation prediction The segmentation image is created by a merge of the model output and the correct data. The color-coded regions in the image correspond to True Positive, False Positive, False Negative, and True Negative regions, respectively, as shown in the upper part of the figure. The upper, middle, and lower rows show the results when the source and target cell type combinations are SH-SY5Y → MCF7, SH-SY5Y → Huh7, and Huh7 → SH-SY5Y, respectively.

A heat map showing the pixel-level discrepancy before and after learning is shown in Fig.6a. In the combination of SH-SY5Y → Huh7 (IoU:0.808 ± 0.061), the high-discrepancy regions tended to be decreased, whereas in the combination of Huh7 → SH-SY5Y (IoU:0.672 ± 0.074), the high-discrepancy regions remained even at the end of learning.

**Figure 6:**
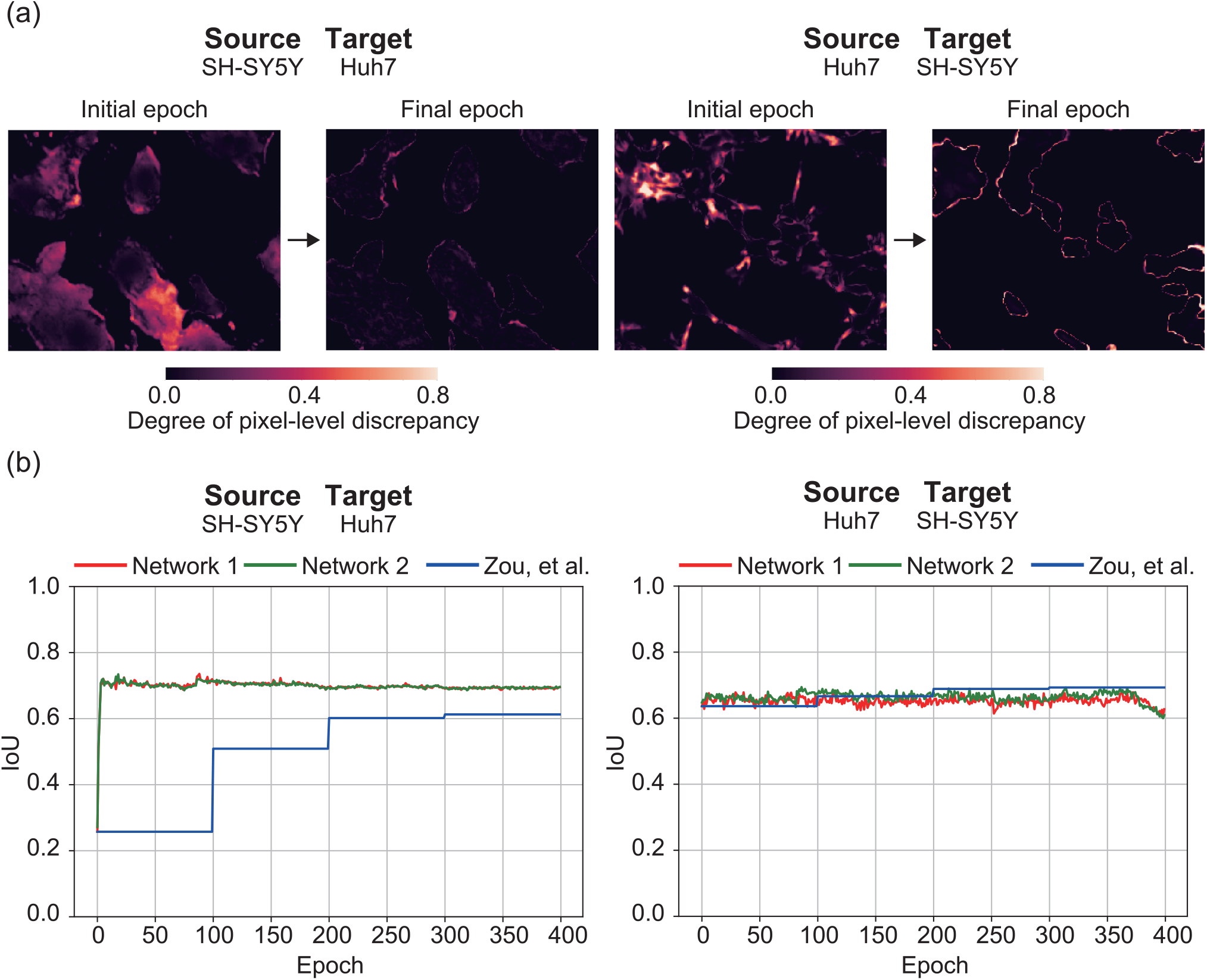
Performance transition in the learning process (a) Transition of the pixel-level discrepancy. Black represents regions with low discrepancy, and red represents regions with high discrepancy. (b) Transition of the IoU for the pseudo-labels. The red and green lines represent the IoUs of the pseudo-labels created from the two networks in the proposed method. The blue line represents the IoUs of the pseudo-labels created using the Zou et al. method [19]. The IoUs in the proposed method are 0.808 ± 0.061 and 0.672 ± 0.074 when the source and target cell type combinations are SH-SY5Y → Huh7 and Huh7 → SH-SY5Y, respectively.

To evaluate the effects of pseudo-label updates on the process of self-learning, we compared the IoU transition of pseudo-labels in the learning process of the proposed method and Zou et al. ‘s method [19] (Fig. 6b). In the SH-SY5Y → Huh7 combination (IoU:0.808 ± 0.061), the IoU of the pseudo-labels in the proposed method improved significantly in the first 10 epochs of learning, whereas the accuracy of the Zou et al. method stopped the accuracy improvement at 300 epochs after a gradual improvement. However, for the combination of Huh7 → SH-SY5Y (IoU: 0.672 ± 0.074), the IoU of the pseudo-labels in the proposed method seemed to decrease gradually during the learning process. This indicates that the proposed method achieved the performance improvement by updating the pseudo-label and model simultaneously, while Zou et al. ‘s method updates pseudo-labels step by step [19].

These results suggested that the simultaneous updating of the pseudo-label and model with consideration of the pixel-level discrepancy improved the segmentation performance by suppressing false negatives in some cell type combinations, such as Huh7 and MCF7 cells. Although the cell types used in the validation of this study were only a subset of the many cell types, our results suggest that we can develop a model that can effectively suppress false-negative errors in unknown cell types by training with the cell images that have protrusion-like structures as a source (Supplementary Fig. S1), such as SH-SY5Y cells, which may have contained the necessary features to constitute cell shape.

## Discussions

To overcome the performance degradation of semantic cell segmentation for an unknown cell type, we developed an unsupervised domain adaptation method based on a cooperative self-learning method that uses two independent models for aligning feature distributions by considering pixel-level discrepancy (Fig. 1). As a result, for the unknown cell types that belong to a different domain from the learned cell type (Supplementary Fig. S4), the proposed method robustly improved the segmentation performance compared with the lower-bound model (Fig. 2, Table 1, Table 2). In addition, for some cell type combinations, the proposed method was not only more efficient and showed higher segmentation performance than conventional approaches, but also showed accuracy comparable to supervised learning models, despite the unsupervised learning (Fig. 3, Fig. 4, Supplementary Tables S2-S5). Furthermore, the proposed method might have improved the accuracy of detecting cellular regions by suppressing false-negative regions through simultaneous updating of pseudo-labels and models considering the pixel-level discrepancy (Fig. 5 and Fig. 6).

Based on the performance evaluation of the supervised U-Net models, the upper-bound model achieved high segmentation accuracy, whereas the lower-bound model showed low segmentation accuracy (Table 1). In comparison, in the results of dimensionality reduction using t-SNE, the segmentation accuracy within the domain of the learned cell type was high, while the segmentation accuracy for cell types in a different domain from the one at the time of learning was low (Supplementary Fig. S4). These results suggested that, as in the previous studies [8], this phenomenon occurred by a domain shift.

Regarding the qualitative performance comparisons, the lower-bound model and conventional models tended to produce false-negative errors that failed to detect cellular regions, noise-like false-negative errors, and false-positive artifacts (Fig. 2, Fig. 3). In contrast, the upper-bound model and the proposed model suppressed these errors. In the target domain, where a domain shift occurs, conventional models tended to make conservative predictions that led to false negatives, whereas the proposed model tended to lead to false positives rather than false negatives. Regarding the quantitative performance comparisons of IoU and DIC, for the combinations with A172, BT-474 and MCF7 cells as the target, the proposed method effectively outperformed the segmentation performance of the upper-bound model and the conventional models (Supplementary Table S2, S3, Fig. 4a, Fig. 4b). Regarding HD95 and ASD, the proposed method performed comparably to the conventional models (Supplementary Table S4, S5, Fig. 4c, Fig. 4d). In addition, the qualitative evaluation of the segmentation results suggested that the performance degradation due to domain shift was caused by an increase in false negatives, and the proposed method, which cooperatively learns pseudo-labels using the pixel-level discrepancy, improved the segmentation performance by suppressing the false negatives (Fig. 5). In the combination of these cell types, the pixel-level discrepancy decreased with learning (Fig. 6a), and the accuracy of the pseudo-label generated based on the discrepancy increased with learning (Fig. 6b). These findings suggest that generating highly accurate pseudo-labels by using the pixel-level discrepancy resulted in high segmentation performance even for unknown cell types.

In contrast, in most combinations of cell types targeting SH-SY5Y cells and A172 cells, the proposed method showed lower segmentation accuracy than the lower-bound model. Cell types other than SH-SY5Y and A172 cells show rounded cell morphology, whereas SH-SY5Y cells and A172 cells have characteristic protrusion-like structures (Supplementary Fig. S1). The qualitative evaluation of the segmentation results indicated that the proposed models trained on the Huh7 cells failed to detect the protrusion-like structures of SH-SY5Y cells, and the false-negative errors seemed to increase (Fig. 5). In addition, the pixel-level discrepancy could not be reduced in these combinations (Fig. 6a), and the accuracy of the pseudo-labels seemed to decrease gradually during the learning process (Fig. 6b). In the case of combinations of cell types with extremely different cell morphologies, the models could not correct the errors by examining the discrepancy, which may have caused the performance degradation as independent models proceeded to learn incorrect labels from each other. Considering these results, we propose that cells with protrusion-like structures contain the necessary features to constitute cell shape, whereas rounded cells contain sufficient features to constitute cell shape. Although the models were only evaluated on a subset of the many cell types, a model trained on cells with characteristic structures as a source using the proposed method can achieve effective unsupervised segmentation, even on unknown cell types. By comparing the dimensionality-reduced feature map using t-SNE and the cell types that show characteristic cell shapes, it might be possible to construct an efficient segmentation technique by selecting cell types that should be prioritized for constructing datasets.

In the field of cell segmentation, handling instance segmentation problems is now common [28]. Instance segmentation technology consists of two parts: a semantic segmentation operation of the region where the cell is located as a region of interest, and a separation operation to distinguish different cells as instances. Our proposed model, CULPICO, conducts only the semantic segmentation and cannot perform the separation operation; however, we can achieve instance segmentation by performing a separation operation using classical image processing techniques, such as the watershed method, on the results of the semantic segmentation predicted by the proposed model. What we want to emphasize here is that, of the two technologies that make up instance segmentation, it is not possible to achieve instance segmentation without the preceding semantic segmentation. One of main contributions of our research is that we have shown that we can perform accurate semantic segmentation for various cell types that are unknown to deep learning models by using the proposed framework, which is the basis of the subsequent cell segmentation. Although the proposed model cannot perform the instance segmentation in an end-to-end manner, we believe that extending the scalability of semantic segmentation will contribute to the development of the cell segmentation field. There is no doubt that extracting individual cell regions using the instance segmentation provides meaningful suggestions for biologists. In addition, there are some cases where extracting cell regions using semantic segmentation rather than instance segmentation itself provides meaningful information for biologists. For example, the computational costs of semantic segmentation models are generally low compared to instance segmentation models; therefore, the analysis time should be shortened for large-scale segmentation analysis on a large amount of microscope image data. In addition, training an instance segmentation model requires training data with each cell labeled with its ID, but a semantic segmentation model only requires training data with each cell region labeled, without requiring a huge number of IDs, so the labor cost borne by biologists is greatly reduced. When the biologists only want to know the area ratio of cell regions in a specific area, building annotation data for an instance segmentation model is an unnecessary burden for them, and, in many cases, the less labor-intensive semantic segmentation model is more appropriate. Moreover, by using our framework, the amount of annotation work required for semantic segmentation models can be greatly reduced, further freeing up time for the biologists. We believe that our model will provide a meaningful experience for biologists who want to spend as much time as possible on biological wet experiments.

## Code availability

The source code of this study is available from https://github.com/funalab/CULPICO.

## Data availability

The datasets including all training and testing datasets during the current study are available from https://drive.google.com/file/d/1knrykZ8aOwNKbOUzQ8a1PvGQodOkY1-V/view?usp=sharing. The datasets are part of a public cell image dataset LIVECell published by Edlund et. al. [22]. Original LIVECell dataset are published under Attribution-NonCommercial 4.0 International (CC BY-NC 4.0) license.

## Acknowledgement

We are grateful to Yuya Kobayashi for valuable discussions. The research was funded by MEXT/JSPS KAKENHI Grant Number JP18H04742 “Resonance Bio” and JST CREST, Japan Grant Number JPMJCR2331 to A.F.

## Author contributions

S.M., S.N., and A.F. designed the conceptual idea and the study. S.M. and T.M. implemented the proposed algorithm and conventional algorithms. T.M. conducted a comprehensive and multifaceted evaluation of the performance of the proposed method. S.M., S.N, Y.T., T.G.Y., T.M., and A.F. wrote the manuscript, with suggestions from the other authors.

