## Supplementary Material for "Cell segmentation without annotation by unsupervised domain adaptation based on cooperative self-learning"

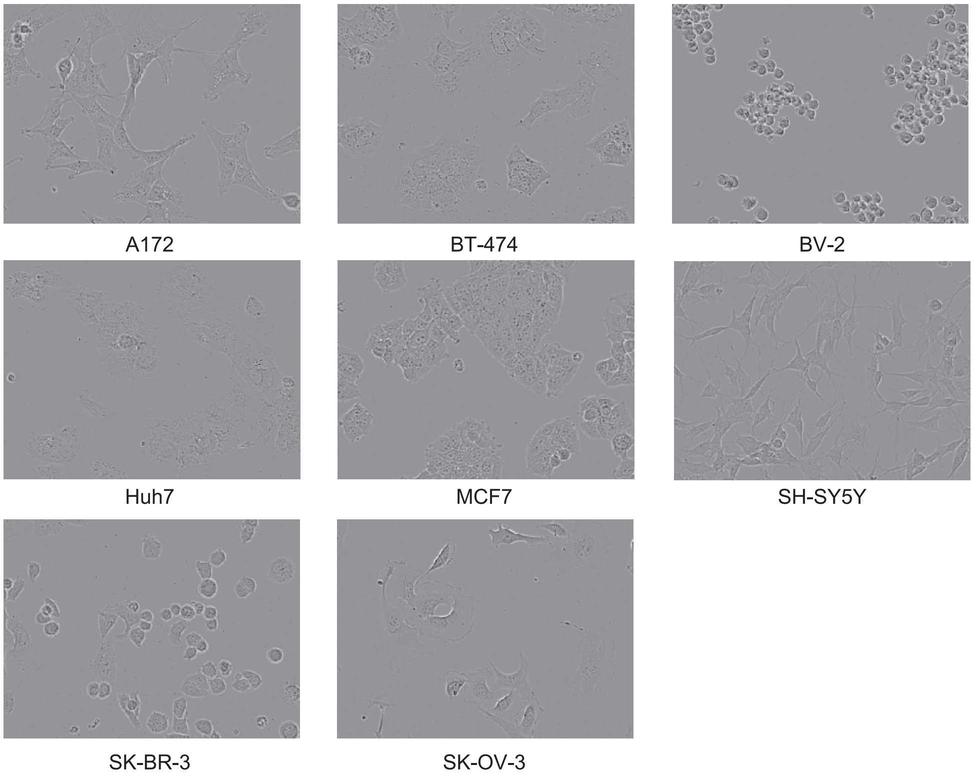


Figure S1: Representative microscopic images included in the dataset

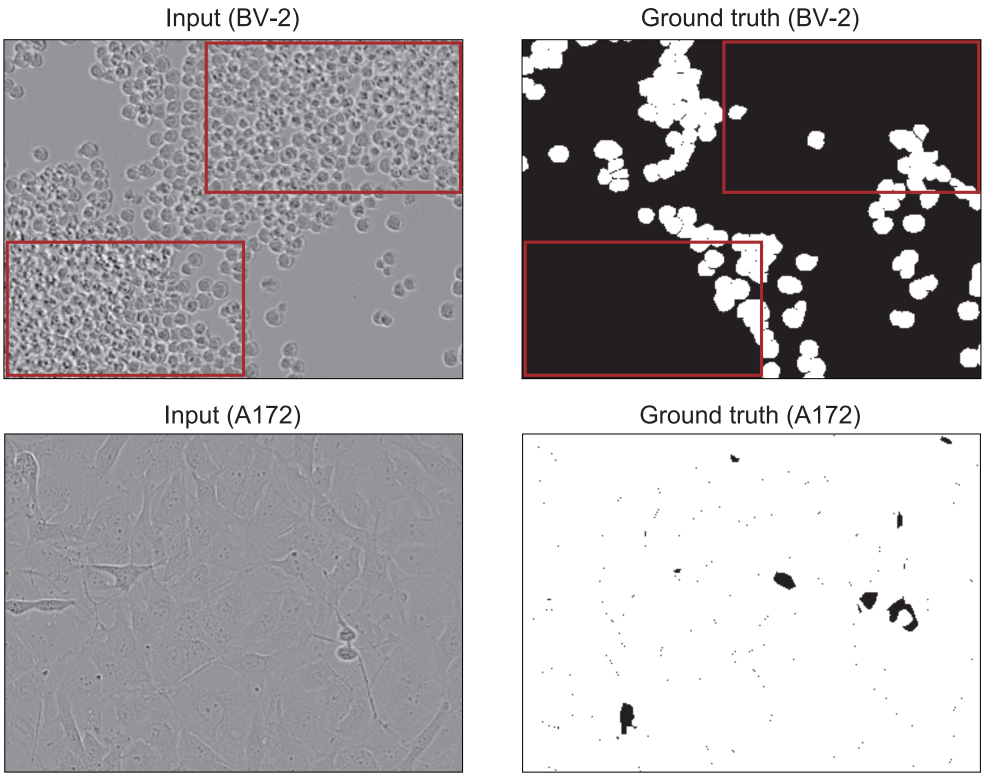


Figure S2: Representative microscopic images excluded from the dataset
In the data of BV-2 cells shown in the upper panels, the areas outlined in red are not annotated. In the data of A172 cells shown in the lower panels, there are regions so dense that the cells cannot be separated.

Table S1: Composition of the number of images in the dataset

| Cell type | Train datasets | Test datasets |
|---|---|---|
| A172 | 366 | 122 |
| BT-474 | 487 | 168 |
| BV-2 | 366 | 126 |
| Huh7 | 400 | 200 |
| MCF7 | 479 | 156 |
| SH-SY5Y | 487 | 165 |
| SK-BR-3 | 527 | 176 |
| SK-OV-3 | 265 | 234 |

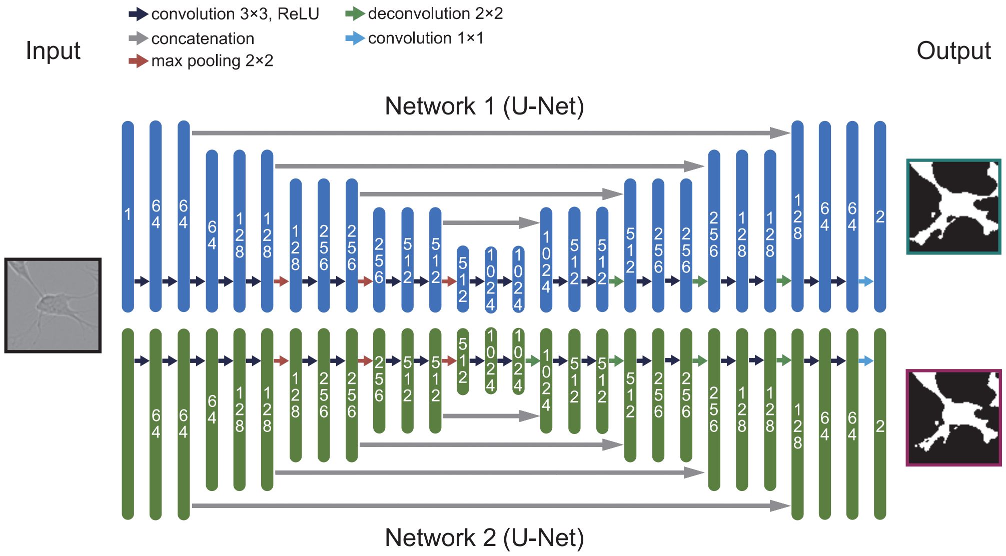


Figure S3: Architecture of the proposed method
The numbers in each layer indicate the number of channels. The color-coded arrows correspond to the processing shown at the top of the figure. When a phase-contrast microscopic image is input to the model, segmentation images are output from two networks.

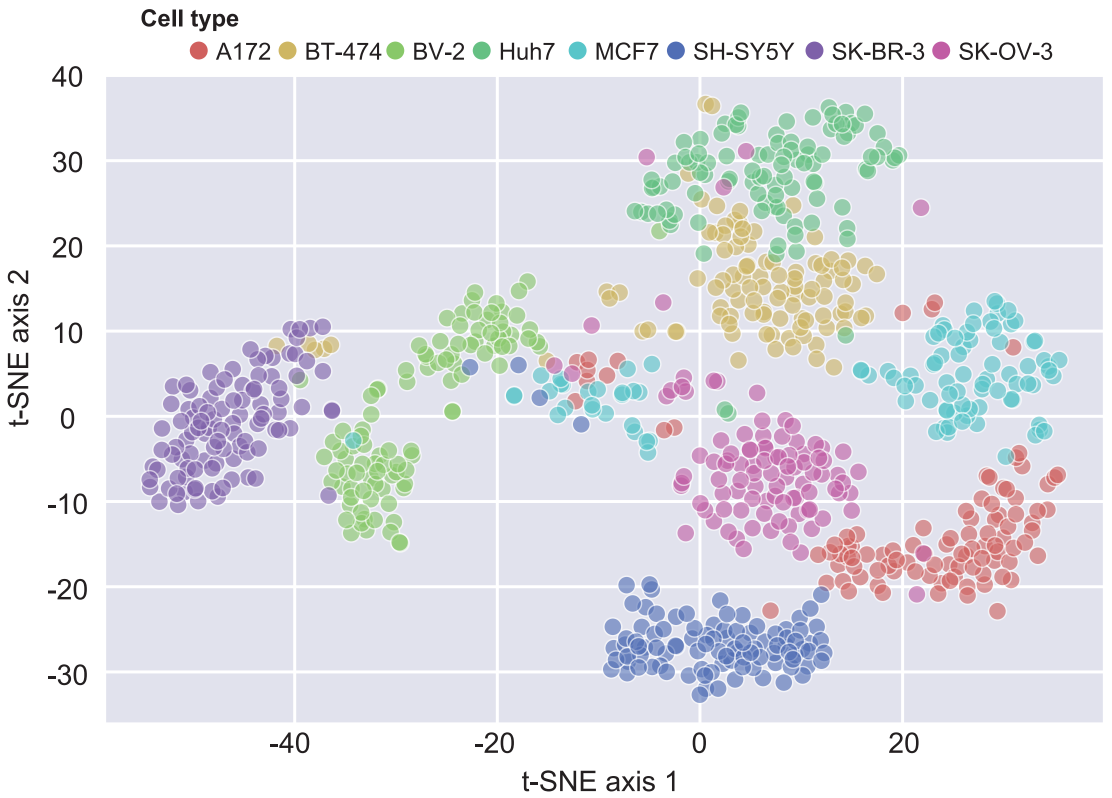


Figure S4: Dimensionality reduction of microscopic cell images into two-dimensional space using t-SNE. The horizontal and vertical axes represent the values in a two-dimensional vector reduced by t-SNE. The plotted points represent the dimension-reduced microscopic images, and the respective colors represent differences in cell types.

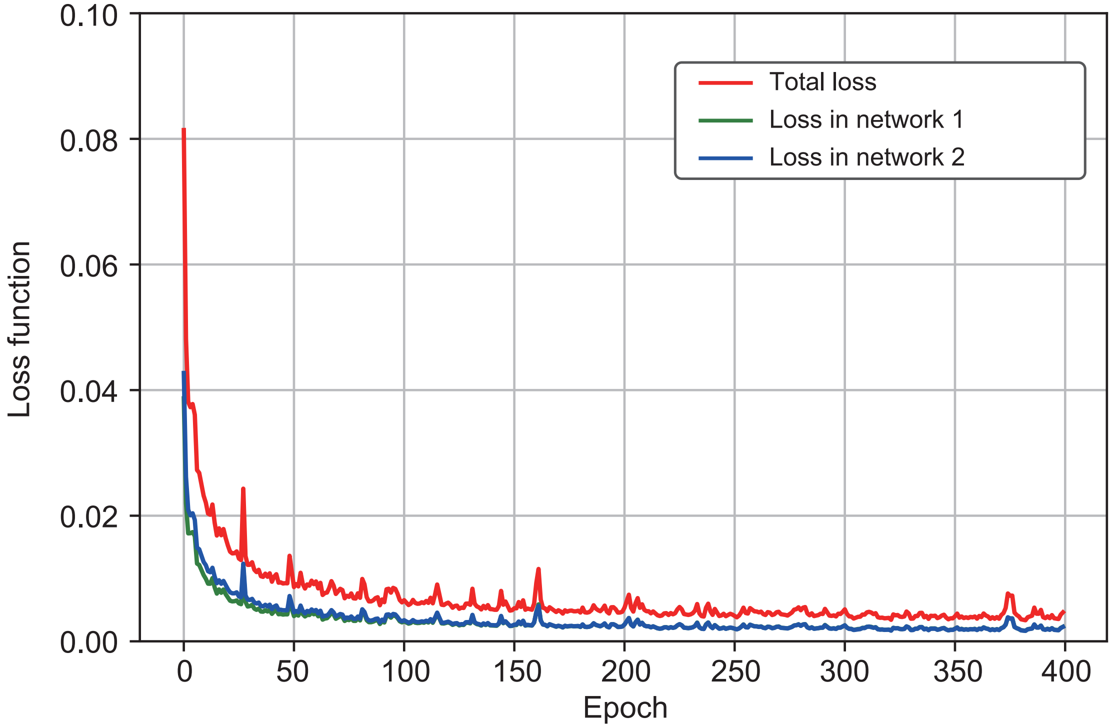


Figure S5: Representative learning curves in the proposed method
The learning curve in the proposed method with SH-SY5Y cells as source and Huh7 cells as target. The horizontal axis represents the number of epochs, and the vertical axis represents the loss value. The red line represents the total loss, and the green and blue lines represent the loss for each network, respectively.

Table S2: Comparison of segmentation performance for the Intersection-over-Union (IoU) metric
Values in the table represent the mean IoU for the test data, and those in parentheses show the standard deviation. Those in bold type represent the model with the highest IoU, except for the upper-bound model. Underlines represent the models with higher IoU than the upper-bound model. The lower-bound model is the model trained with only source data. The upper-bound model is the model trained with only target data. To save space, the names of the source and target cell lines have been abbreviated as follows: BT-474 to BT47, BV-2 to BV2, SH-SY5Y to SHSY, SK-BR-3 to SKBR, and SK-OV-3 to SKOV.

| Source | Target | Lower bound | Saito | Zou | Na | Wang | Ours | Upper bound | Source | Target | Lower bound | Saito | Zou | Na | Wang | Ours | Upper bound |
|---|---|---|---|---|---|---|---|---|---|---|---|---|---|---|---|---|---|
| A172 | BT47 | 0.790 (0.078) | 0.417 (0.177) | 0.804 (0.094) | **0.820** (0.081) | 0.624 (0.076) | 0.784 (0.074) | 0.817 (0.055) | MCF7 | A172 | 0.902 (0.027) | 0.815 (0.037) | 0.900 (0.029) | **0.907** (0.032) | 0.563 (0.161) | 0.820 (0.045) | 0.878 (0.029) |
| A172 | BV2 | 0.804 (0.083) | 0.799 (0.084) | 0.784 (0.082) | **0.819** (0.089) | 0.548 (0.086) | 0.518 (0.079) | 0.842 (0.064) | MCF7 | BT47 | 0.777 (0.072) | 0.794 (0.070) | 0.797 (0.085) | **0.826** (0.068) | 0.606 (0.070) | 0.819 (0.080) | 0.817 (0.055) |
| A172 | Huh7 | 0.664 (0.063) | 0.343 (0.152) | 0.844 (0.058) | **0.848** (0.058) | 0.682 (0.059) | 0.762 (0.065) | 0.867 (0.054) | MCF7 | BV2 | 0.813 (0.077) | 0.757 (0.070) | 0.826 (0.085) | **0.835** (0.072) | 0.537 (0.086) | 0.786 (0.068) | 0.842 (0.064) |
| A172 | MCF7 | 0.858 (0.067) | 0.860 (0.070) | **0.866** (0.069) | 0.864 (0.070) | 0.755 (0.088) | 0.857 (0.068) | 0.847 (0.063) | MCF7 | Huh7 | 0.711 (0.060) | 0.715 (0.058) | 0.751 (0.059) | **0.822** (0.056) | 0.533 (0.131) | 0.806 (0.060) | 0.867 (0.054) |
| A172 | SHSY | 0.721 (0.066) | 0.725 (0.075) | **0.762** (0.066) | 0.756 (0.067) | 0.585 (0.074) | 0.617 (0.082) | 0.789 (0.052) | MCF7 | SHSY | 0.739 (0.062) | 0.716 (0.064) | 0.744 (0.066) | **0.759** (0.063) | 0.537 (0.083) | 0.630 (0.080) | 0.789 (0.052) |
| A172 | SKBR | 0.865 (0.024) | 0.879 (0.033) | **0.891** (0.032) | 0.888 (0.042) | 0.677 (0.063) | 0.837 (0.052) | 0.903 (0.035) | MCF7 | SKBR | 0.866 (0.033) | 0.837 (0.028) | 0.878 (0.027) | **0.885** (0.035) | 0.607 (0.094) | 0.872 (0.029) | 0.903 (0.035) |
| A172 | SKOV | 0.486 (0.104) | 0.844 (0.052) | **0.887** (0.044) | 0.863 (0.049) | 0.753 (0.083) | 0.884 (0.051) | 0.917 (0.030) | MCF7 | SKOV | 0.660 (0.098) | 0.763 (0.050) | 0.653 (0.080) | **0.841** (0.053) | 0.557 (0.099) | 0.632 (0.071) | 0.917 (0.030) |
| BT47 | A172 | **0.910** (0.030) | 0.707 (0.053) | 0.894 (0.036) | 0.888 (0.028) | 0.661 (0.077) | 0.852 (0.038) | 0.878 (0.029) | SHSY | A172 | 0.493 (0.095) | 0.764 (0.036) | 0.812 (0.049) | 0.895 (0.031) | 0.677 (0.101) | **0.907** (0.028) | 0.878 (0.029) |
| BT47 | BV2 | **0.827** (0.072) | 0.692 (0.092) | 0.802 (0.073) | 0.824 (0.068) | 0.513 (0.086) | 0.779 (0.068) | 0.842 (0.064) | SHSY | BT47 | 0.375 (0.184) | 0.686 (0.074) | 0.769 (0.073) | 0.813 (0.079) | 0.601 (0.065) | **0.814** (0.078) | 0.817 (0.055) |
| BT47 | Huh7 | 0.795 (0.057) | 0.685 (0.071) | 0.845 (0.052) | 0.847 (0.055) | 0.650 (0.059) | **0.861** (0.058) | 0.867 (0.054) | SHSY | BV2 | 0.838 (0.061) | 0.786 (0.060) | **0.840** (0.064) | 0.838 (0.065) | 0.496 (0.083) | 0.827 (0.065) | 0.842 (0.064) |
| BT47 | MCF7 | 0.810 (0.089) | 0.822 (0.076) | 0.858 (0.065) | **0.863** (0.067) | 0.724 (0.087) | 0.861 (0.067) | 0.847 (0.063) | SHSY | Huh7 | 0.156 (0.143) | 0.688 (0.059) | 0.623 (0.110) | 0.800 (0.052) | 0.553 (0.081) | **0.808** (0.061) | 0.867 (0.054) |
| BT47 | SHSY | **0.759** (0.064) | 0.682 (0.062) | 0.707 (0.069) | 0.724 (0.068) | 0.548 (0.076) | 0.712 (0.078) | 0.789 (0.052) | SHSY | MCF7 | 0.571 (0.167) | 0.838 (0.061) | 0.822 (0.082) | 0.862 (0.068) | 0.715 (0.095) | **0.863** (0.070) | 0.847 (0.063) |
| BT47 | SKBR | 0.887 (0.036) | 0.797 (0.027) | 0.866 (0.031) | **0.894** (0.035) | 0.554 (0.084) | 0.891 (0.031) | 0.903 (0.035) | SHSY | SKBR | 0.789 (0.053) | 0.809 (0.034) | 0.840 (0.041) | 0.866 (0.035) | 0.604 (0.076) | **0.881** (0.029) | 0.903 (0.035) |
| BT47 | SKOV | 0.847 (0.048) | 0.613 (0.049) | 0.873 (0.043) | 0.805 (0.075) | 0.471 (0.072) | **0.875** (0.054) | 0.917 (0.030) | SHSY | SKOV | 0.575 (0.085) | 0.674 (0.039) | 0.684 (0.058) | 0.789 (0.043) | 0.479 (0.064) | **0.813** (0.060) | 0.917 (0.030) |
| BV2 | A172 | 0.161 (0.153) | 0.579 (0.060) | 0.324 (0.175) | 0.833 (0.036) | 0.094 (0.090) | **0.860** (0.036) | 0.878 (0.029) | SKBR | A172 | **0.918** (0.026) | 0.775 (0.213) | 0.912 (0.048) | 0.918 (0.030) | 0.777 (0.070) | 0.912 (0.028) | 0.878 (0.029) |
| BV2 | BT47 | 0.212 (0.237) | 0.682 (0.092) | 0.366 (0.235) | 0.777 (0.080) | 0.371 (0.099) | **0.814** (0.079) | 0.817 (0.055) | SKBR | BT47 | **0.826** (0.080) | 0.783 (0.082) | 0.822 (0.080) | 0.821 (0.079) | 0.634 (0.076) | 0.824 (0.079) | 0.817 (0.055) |
| BV2 | Huh7 | 0.048 (0.044) | 0.563 (0.050) | 0.116 (0.091) | **0.751** (0.060) | 0.217 (0.050) | 0.579 (0.055) | 0.867 (0.054) | SKBR | BV2 | 0.834 (0.076) | 0.814 (0.084) | **0.834** (0.068) | 0.826 (0.090) | 0.602 (0.077) | 0.826 (0.088) | 0.842 (0.064) |
| BV2 | MCF7 | 0.291 (0.214) | 0.733 (0.081) | 0.498 (0.203) | 0.843 (0.063) | 0.515 (0.104) | **0.854** (0.069) | 0.847 (0.063) | SKBR | Huh7 | **0.864** (0.055) | 0.742 (0.086) | 0.842 (0.059) | 0.841 (0.057) | 0.635 (0.068) | 0.850 (0.059) | 0.867 (0.054) |
| BV2 | SHSY | 0.410 (0.112) | 0.686 (0.057) | 0.523 (0.092) | **0.739** (0.066) | 0.313 (0.119) | 0.692 (0.071) | 0.789 (0.052) | SKBR | MCF7 | 0.863 (0.070) | 0.863 (0.069) | 0.862 (0.071) | **0.865** (0.070) | 0.781 (0.078) | 0.859 (0.071) | 0.847 (0.063) |
| BV2 | SKBR | 0.745 (0.063) | 0.690 (0.083) | 0.719 (0.075) | 0.790 (0.052) | 0.395 (0.083) | **0.875** (0.031) | 0.903 (0.035) | SKBR | SHSY | 0.693 (0.085) | 0.717 (0.077) | 0.732 (0.077) | **0.770** (0.067) | 0.597 (0.069) | 0.719 (0.079) | 0.789 (0.052) |
| BV2 | SKOV | 0.092 (0.054) | 0.480 (0.061) | 0.168 (0.076) | **0.700** (0.047) | 0.072 (0.033) | 0.247 (0.060) | 0.917 (0.030) | SKBR | SKOV | **0.895** (0.037) | 0.840 (0.052) | 0.753 (0.069) | 0.858 (0.044) | 0.737 (0.076) | 0.569 (0.064) | 0.917 (0.030) |
| Huh7 | A172 | **0.915** (0.027) | 0.494 (0.182) | 0.887 (0.038) | 0.871 (0.040) | 0.710 (0.120) | 0.909 (0.030) | 0.878 (0.029) | SKOV | A172 | 0.917 (0.034) | 0.913 (0.033) | 0.883 (0.079) | **0.917** (0.032) | 0.758 (0.091) | 0.911 (0.037) | 0.878 (0.029) |
| Huh7 | BT47 | 0.771 (0.099) | 0.735 (0.178) | 0.782 (0.093) | **0.822** (0.075) | 0.550 (0.117) | 0.820 (0.078) | 0.817 (0.055) | SKOV | BT47 | 0.785 (0.098) | 0.553 (0.138) | 0.751 (0.112) | 0.801 (0.089) | 0.586 (0.087) | **0.806** (0.080) | 0.817 (0.055) |
| Huh7 | BV2 | **0.795** (0.110) | 0.176 (0.156) | 0.726 (0.167) | 0.792 (0.071) | 0.371 (0.115) | 0.686 (0.142) | 0.842 (0.064) | SKOV | BV2 | 0.787 (0.089) | 0.784 (0.096) | 0.795 (0.090) | 0.787 (0.092) | 0.513 (0.102) | **0.800** (0.092) | 0.842 (0.064) |
| Huh7 | MCF7 | 0.731 (0.192) | 0.606 (0.253) | 0.730 (0.193) | **0.861** (0.069) | 0.651 (0.116) | 0.856 (0.072) | 0.847 (0.063) | SKOV | Huh7 | 0.837 (0.070) | 0.755 (0.092) | 0.840 (0.067) | **0.853** (0.060) | 0.678 (0.063) | 0.844 (0.061) | 0.867 (0.054) |
| Huh7 | SHSY | **0.738** (0.054) | 0.520 (0.134) | 0.677 (0.068) | 0.699 (0.065) | 0.492 (0.099) | 0.672 (0.074) | 0.789 (0.052) | SKOV | MCF7 | 0.858 (0.074) | 0.283 (0.168) | 0.851 (0.074) | **0.860** (0.073) | 0.724 (0.107) | 0.854 (0.071) | 0.847 (0.063) |
| Huh7 | SKBR | **0.889** (0.029) | 0.752 (0.042) | 0.873 (0.033) | 0.885 (0.029) | 0.469 (0.129) | 0.879 (0.041) | 0.903 (0.035) | SKOV | SHSY | **0.765** (0.071) | 0.578 (0.149) | 0.740 (0.078) | 0.743 (0.068) | 0.560 (0.081) | 0.711 (0.075) | 0.789 (0.052) |
| Huh7 | SKOV | 0.825 (0.084) | 0.682 (0.071) | 0.873 (0.054) | 0.744 (0.107) | 0.693 (0.100) | **0.912** (0.031) | 0.917 (0.030) | SKOV | SKBR | 0.876 (0.051) | 0.411 (0.056) | 0.865 (0.052) | **0.890** (0.040) | 0.650 (0.077) | 0.868 (0.048) | 0.903 (0.035) |

Table S3: Comparison of segmentation performance for the Dice coefficient
Values in the table represent the mean Dice coefficient for the test data, and those in parentheses show the standard deviation. Those in bold type represent the model with the highest Dice coefficient, except for the upper-bound model. Underlines represent the models with higher Dice coefficient than the upper-bound model. The lower-bound model is the model trained with only source data. The upper-bound model is the model trained with only target data. To save space, the names of the source and target cell lines have been abbreviated as follows: BT-474 to BT47, BV-2 to BV2, SH-SY5Y to SHSY, SK-BR-3 to SKBR, and SK-OV-3 to SKOV.

| Source | Target | Lower bound | Saito | Zou | Na | Wang | Ours | Upper bound | Source | Target | Lower bound | Saito | Zou | Na | Wang | Ours | Upper bound |
|---|---|---|---|---|---|---|---|---|---|---|---|---|---|---|---|---|---|
| A172 | BT47 | 0.881 (0.052) | 0.566 (0.182) | 0.888 (0.063) | **0.899** (0.052) | 0.766 (0.060) | 0.877 (0.049) | 0.898 (0.034) | MCF7 | A172 | 0.948 (0.015) | 0.898 (0.023) | 0.947 (0.016) | **0.951** (0.017) | 0.706 (0.144) | 0.901 (0.028) | 0.935 (0.017) |
| A172 | BV2 | 0.889 (0.058) | 0.885 (0.059) | 0.877 (0.058) | **0.898** (0.061) | 0.704 (0.075) | 0.679 (0.070) | 0.913 (0.042) | MCF7 | BT47 | 0.873 (0.047) | 0.883 (0.046) | 0.884 (0.056) | **0.903** (0.043) | 0.752 (0.055) | 0.898 (0.051) | 0.898 (0.034) |
| A172 | Huh7 | 0.796 (0.047) | 0.493 (0.163) | 0.915 (0.036) | **0.917** (0.036) | 0.809 (0.043) | 0.864 (0.044) | 0.928 (0.033) | MCF7 | BV2 | 0.895 (0.050) | 0.860 (0.048) | 0.902 (0.057) | **0.908** (0.046) | 0.695 (0.076) | 0.878 (0.046) | 0.913 (0.042) |
| A172 | MCF7 | 0.922 (0.040) | 0.923 (0.042) | **0.926** (0.042) | 0.926 (0.042) | 0.858 (0.059) | 0.921 (0.041) | 0.916 (0.038) | MCF7 | Huh7 | 0.830 (0.042) | 0.832 (0.041) | 0.856 (0.040) | **0.901** (0.036) | 0.685 (0.118) | 0.891 (0.039) | 0.928 (0.033) |
| A172 | SHSY | 0.836 (0.045) | 0.838 (0.051) | **0.863** (0.043) | 0.859 (0.044) | 0.736 (0.058) | 0.760 (0.064) | 0.881 (0.033) | MCF7 | SHSY | 0.849 (0.041) | 0.833 (0.044) | 0.851 (0.043) | **0.862** (0.041) | 0.695 (0.070) | 0.770 (0.061) | 0.881 (0.033) |
| A172 | SKBR | 0.927 (0.014) | 0.935 (0.020) | **0.942** (0.019) | 0.940 (0.024) | 0.806 (0.045) | 0.910 (0.032) | 0.949 (0.020) | MCF7 | SKBR | 0.928 (0.019) | 0.911 (0.017) | 0.935 (0.016) | **0.939** (0.020) | 0.751 (0.075) | 0.931 (0.017) | 0.949 (0.020) |
| A172 | SKOV | 0.648 (0.094) | 0.914 (0.032) | **0.939** (0.025) | 0.926 (0.029) | 0.856 (0.054) | 0.938 (0.029) | 0.957 (0.017) | MCF7 | SKOV | 0.791 (0.073) | 0.865 (0.033) | 0.787 (0.059) | **0.913** (0.033) | 0.710 (0.090) | 0.772 (0.054) | 0.957 (0.017) |
| BT47 | A172 | **0.953** (0.017) | 0.827 (0.036) | 0.943 (0.021) | 0.940 (0.015) | 0.793 (0.055) | 0.920 (0.022) | 0.935 (0.017) | SHSY | A172 | 0.655 (0.086) | 0.866 (0.023) | 0.895 (0.030) | 0.944 (0.017) | 0.803 (0.075) | **0.951** (0.016) | 0.935 (0.017) |
| BT47 | BV2 | **0.903** (0.046) | 0.814 (0.067) | 0.888 (0.048) | 0.902 (0.044) | 0.674 (0.080) | 0.874 (0.045) | 0.913 (0.042) | SHSY | BT47 | 0.522 (0.171) | 0.811 (0.053) | 0.868 (0.047) | 0.895 (0.051) | 0.749 (0.052) | **0.895** (0.050) | 0.898 (0.034) |
| BT47 | Huh7 | 0.885 (0.036) | 0.811 (0.050) | 0.915 (0.032) | 0.916 (0.034) | 0.786 (0.044) | **0.924** (0.036) | 0.928 (0.033) | SHSY | BV2 | 0.911 (0.037) | 0.879 (0.039) | **0.912** (0.040) | 0.910 (0.041) | 0.658 (0.079) | 0.904 (0.041) | 0.913 (0.042) |
| BT47 | MCF7 | 0.893 (0.057) | 0.900 (0.047) | 0.922 (0.039) | **0.925** (0.041) | 0.837 (0.060) | 0.924 (0.040) | 0.916 (0.038) | SHSY | Huh7 | 0.248 (0.183) | 0.814 (0.043) | 0.762 (0.084) | 0.888 (0.034) | 0.708 (0.072) | **0.893** (0.039) | 0.928 (0.033) |
| BT47 | SHSY | **0.862** (0.042) | 0.809 (0.044) | 0.827 (0.047) | 0.838 (0.045) | 0.705 (0.061) | 0.829 (0.055) | 0.881 (0.033) | SHSY | MCF7 | 0.712 (0.141) | 0.911 (0.037) | 0.900 (0.052) | 0.924 (0.041) | 0.830 (0.066) | **0.925** (0.042) | 0.916 (0.038) |
| BT47 | SKBR | 0.940 (0.021) | 0.887 (0.017) | 0.928 (0.018) | **0.943** (0.020) | 0.709 (0.072) | 0.942 (0.018) | 0.949 (0.020) | SHSY | SKBR | 0.881 (0.033) | 0.894 (0.021) | 0.913 (0.025) | 0.928 (0.021) | 0.751 (0.060) | **0.937** (0.017) | 0.949 (0.020) |
| BT47 | SKOV | 0.917 (0.029) | 0.759 (0.038) | 0.931 (0.025) | 0.890 (0.048) | 0.637 (0.067) | **0.933** (0.032) | 0.957 (0.017) | SHSY | SKOV | 0.726 (0.069) | 0.804 (0.028) | 0.811 (0.041) | 0.882 (0.027) | 0.645 (0.058) | **0.896** (0.037) | 0.957 (0.017) |
| BV2 | A172 | 0.252 (0.192) | 0.731 (0.048) | 0.466 (0.184) | 0.908 (0.022) | 0.161 (0.141) | **0.925** (0.021) | 0.935 (0.017) | SKBR | A172 | **0.957** (0.014) | 0.851 (0.185) | 0.953 (0.030) | 0.957 (0.017) | 0.873 (0.045) | 0.954 (0.016) | 0.935 (0.017) |
| BV2 | BT47 | 0.300 (0.258) | 0.807 (0.067) | 0.497 (0.226) | 0.872 (0.053) | 0.533 (0.108) | **0.896** (0.051) | 0.898 (0.034) | SKBR | BT47 | **0.903** (0.051) | 0.876 (0.055) | 0.900 (0.051) | 0.900 (0.051) | 0.773 (0.060) | 0.901 (0.050) | 0.898 (0.034) |
| BV2 | Huh7 | 0.088 (0.074) | 0.719 (0.042) | 0.197 (0.132) | **0.857** (0.040) | 0.354 (0.066) | 0.732 (0.045) | 0.928 (0.033) | SKBR | BV2 | 0.907 (0.050) | 0.895 (0.059) | **0.908** (0.043) | 0.902 (0.061) | 0.749 (0.062) | 0.902 (0.060) | 0.913 (0.042) |
| BV2 | MCF7 | 0.412 (0.238) | 0.843 (0.058) | 0.639 (0.190) | 0.914 (0.038) | 0.674 (0.090) | **0.919** (0.041) | 0.916 (0.038) | SKBR | Huh7 | **0.926** (0.034) | 0.849 (0.059) | 0.913 (0.037) | 0.912 (0.035) | 0.775 (0.054) | 0.918 (0.037) | 0.928 (0.033) |
| BV2 | SHSY | 0.574 (0.096) | 0.812 (0.040) | 0.682 (0.073) | **0.848** (0.044) | 0.464 (0.147) | 0.816 (0.050) | 0.881 (0.033) | SKBR | MCF7 | 0.925 (0.042) | 0.925 (0.041) | 0.924 (0.043) | **0.926** (0.042) | 0.875 (0.050) | 0.923 (0.042) | 0.916 (0.038) |
| BV2 | SKBR | 0.853 (0.041) | 0.813 (0.059) | 0.835 (0.050) | 0.882 (0.033) | 0.562 (0.088) | **0.933** (0.018) | 0.949 (0.020) | SKBR | SHSY | 0.816 (0.058) | 0.833 (0.053) | 0.843 (0.052) | **0.869** (0.044) | 0.745 (0.054) | 0.834 (0.055) | 0.881 (0.033) |
| BV2 | SKOV | 0.164 (0.084) | 0.646 (0.056) | 0.280 (0.109) | **0.823** (0.033) | 0.132 (0.056) | 0.392 (0.076) | 0.957 (0.017) | SKBR | SKOV | **0.944** (0.021) | 0.912 (0.031) | 0.857 (0.046) | 0.923 (0.026) | 0.846 (0.050) | 0.724 (0.052) | 0.957 (0.017) |
| Huh7 | A172 | **0.955** (0.015) | 0.641 (0.169) | 0.940 (0.022) | 0.930 (0.023) | 0.824 (0.088) | 0.952 (0.017) | 0.935 (0.017) | SKOV | A172 | 0.956 (0.018) | 0.954 (0.018) | 0.936 (0.052) | **0.957** (0.017) | 0.859 (0.061) | 0.953 (0.020) | 0.935 (0.017) |
| Huh7 | BT47 | 0.867 (0.066) | 0.831 (0.162) | 0.874 (0.061) | **0.901** (0.047) | 0.702 (0.106) | 0.899 (0.050) | 0.898 (0.034) | SKOV | BT47 | 0.876 (0.067) | 0.702 (0.118) | 0.853 (0.081) | 0.887 (0.059) | 0.735 (0.071) | **0.890** (0.052) | 0.898 (0.034) |
| Huh7 | BV2 | 0.881 (0.075) | 0.270 (0.218) | 0.828 (0.142) | **0.882** (0.047) | 0.530 (0.131) | 0.804 (0.115) | 0.913 (0.042) | SKOV | BV2 | 0.878 (0.061) | 0.875 (0.068) | 0.882 (0.062) | 0.878 (0.064) | 0.672 (0.096) | **0.886** (0.063) | 0.913 (0.042) |
| Huh7 | MCF7 | 0.828 (0.151) | 0.720 (0.224) | 0.828 (0.151) | **0.924** (0.042) | 0.782 (0.093) | 0.921 (0.044) | 0.916 (0.038) | SKOV | Huh7 | 0.910 (0.045) | 0.857 (0.064) | 0.911 (0.043) | **0.920** (0.037) | 0.806 (0.047) | 0.914 (0.038) | 0.928 (0.033) |
| Huh7 | SHSY | **0.848** (0.036) | 0.673 (0.120) | 0.805 (0.048) | 0.821 (0.045) | 0.653 (0.091) | 0.801 (0.052) | 0.881 (0.033) | SKOV | MCF7 | 0.922 (0.046) | 0.418 (0.179) | 0.918 (0.045) | **0.923** (0.045) | 0.835 (0.077) | 0.920 (0.043) | 0.916 (0.038) |
| Huh7 | SKBR | **0.941** (0.017) | 0.858 (0.028) | 0.932 (0.019) | 0.939 (0.017) | 0.627 (0.127) | 0.935 (0.024) | 0.949 (0.020) | SKOV | SHSY | **0.865** (0.047) | 0.720 (0.126) | 0.848 (0.053) | 0.851 (0.046) | 0.714 (0.065) | 0.829 (0.053) | 0.881 (0.033) |
| Huh7 | SKOV | 0.901 (0.053) | 0.809 (0.051) | 0.931 (0.032) | 0.849 (0.076) | 0.815 (0.071) | **0.954** (0.017) | 0.957 (0.017) | SKOV | SKBR | 0.933 (0.030) | 0.581 (0.056) | 0.927 (0.031) | **0.942** (0.023) | 0.785 (0.057) | 0.929 (0.028) | 0.949 (0.020) |

Table S4: Comparison of segmentation performance for the 95% Hausdorff distance (HD95)
Values in the table represent the mean HD95 for the test data, and those in parentheses show the standard deviation. Those in bold type represent the model with the lowest HD95, except for the upper-bound model. Underlines represent the models with lower HD95 than the upper-bound model. The lower-bound model is the model trained with only source data. The upper-bound model is the model trained with only target data. To save space, the names of the source and target cell lines have been abbreviated as follows: BT-474 to BT47, BV-2 to BV2, SH-SY5Y to SHSY, SK-BR-3 to SKBR, and SK-OV-3 to SKOV.

| Source | Target | Lower bound | Saito | Zou | Na | Wang | Ours | Upper bound | Source | Target | Lower bound | Saito | Zou | Na | Wang | Ours | Upper bound |
|---|---|---|---|---|---|---|---|---|---|---|---|---|---|---|---|---|---|
| A172 | BT47 | **22.49** (9.498) | 38.85 (8.177) | 24.15 (10.89) | 23.25 (10.33) | 27.07 (11.21) | 24.77 (10.56) | 17.91 (7.316) | MCF7 | A172 | 15.78 (11.03) | **12.61** (4.194) | 14.07 (8.928) | 16.68 (11.20) | 31.72 (17.63) | 12.66 (5.449) | 12.35 (6.537) |
| A172 | BV2 | 8.127 (5.469) | 9.586 (6.494) | 12.86 (6.543) | **7.472** (5.799) | 17.04 (6.620) | 32.22 (26.36) | 5.574 (5.844) | MCF7 | BT47 | **18.11** (7.494) | 21.61 (8.676) | 23.26 (9.987) | 21.31 (9.096) | 22.66 (7.470) | 22.80 (10.39) | 17.91 (7.316) |
| A172 | Huh7 | **23.01** (11.26) | 77.48 (20.74) | 27.50 (14.68) | 31.39 (15.67) | 39.96 (16.41) | 41.95 (16.05) | 25.46 (14.63) | MCF7 | BV2 | 8.047 (7.168) | 9.175 (6.199) | 8.848 (7.728) | **8.041** (6.020) | 16.76 (7.293) | 9.080 (4.444) | 5.574 (5.844) |
| A172 | MCF7 | **34.62** (28.90) | 36.66 (32.00) | 36.21 (32.02) | 36.24 (31.87) | 37.66 (27.47) | 36.28 (32.00) | 25.56 (13.18) | MCF7 | Huh7 | **22.20** (12.37) | 30.82 (14.95) | 24.88 (13.98) | 24.65 (13.44) | 42.71 (21.94) | 29.40 (14.72) | 25.46 (14.63) |
| A172 | SHSY | **9.837** (2.229) | 15.07 (7.690) | 10.84 (4.245) | 11.10 (6.180) | 26.03 (13.69) | 41.30 (28.10) | 6.468 (2.137) | MCF7 | SHSY | **10.42** (4.105) | 12.91 (7.971) | 10.44 (3.248) | 10.45 (6.167) | 28.48 (15.46) | 28.98 (16.24) | 6.468 (2.137) |
| A172 | SKBR | **7.100** (3.068) | 9.484 (4.994) | 8.216 (4.649) | 8.842 (4.814) | 18.22 (7.209) | 11.88 (5.730) | 5.810 (3.341) | MCF7 | SKBR | **6.255** (2.613) | 7.862 (3.011) | 7.014 (3.208) | 7.412 (3.698) | 18.13 (5.412) | 7.408 (3.299) | 5.810 (3.341) |
| A172 | SKOV | **16.91** (3.256) | 22.61 (14.38) | 18.94 (17.57) | 30.68 (25.54) | 37.70 (22.90) | 40.26 (43.50) | 26.63 (31.72) | MCF7 | SKOV | 16.62 (4.184) | **13.55** (3.167) | 14.19 (2.713) | 19.31 (9.644) | 24.44 (5.600) | 16.04 (2.990) | 26.63 (31.72) |
| BT47 | A172 | 14.46 (10.24) | **12.74** (2.649) | 13.80 (8.532) | 17.61 (11.72) | 20.22 (5.278) | 13.12 (6.849) | 12.35 (6.537) | SHSY | A172 | 18.30 (6.287) | **10.66** (2.108) | 10.67 (2.676) | 13.08 (5.674) | 21.69 (8.234) | 17.29 (12.89) | 12.35 (6.537) |
| BT47 | BV2 | **7.328** (4.753) | 21.73 (14.17) | 9.940 (6.180) | 9.213 (6.087) | 17.61 (9.889) | 17.90 (11.14) | 5.574 (5.844) | SHSY | BT47 | 34.73 (11.76) | 32.56 (6.752) | **16.11** (5.934) | 16.11 (5.905) | 22.74 (8.240) | 21.76 (9.828) | 17.91 (7.316) |
| BT47 | Huh7 | 25.30 (13.72) | **23.19** (13.42) | 24.67 (13.98) | 27.73 (14.60) | 30.79 (15.15) | 30.29 (15.85) | 25.46 (14.63) | SHSY | BV2 | **5.930** (4.755) | 6.537 (4.918) | 6.679 (5.226) | 7.158 (5.528) | 19.05 (14.16) | 7.810 (5.468) | 5.574 (5.844) |
| BT47 | MCF7 | **22.10** (10.08) | 22.94 (12.32) | 28.13 (15.87) | 29.93 (19.04) | 23.31 (9.651) | 34.19 (29.34) | 25.56 (13.18) | SHSY | Huh7 | 80.44 (25.32) | 31.06 (16.26) | **24.20** (13.61) | 25.79 (14.80) | 31.73 (15.83) | 27.77 (14.19) | 25.46 (14.63) |
| BT47 | SHSY | **9.298** (5.517) | 10.71 (2.295) | 12.22 (5.181) | 12.68 (3.142) | 21.65 (6.230) | 19.59 (10.79) | 6.468 (2.137) | SHSY | MCF7 | 24.28 (10.99) | 22.52 (11.90) | **21.51** (9.496) | 35.32 (31.08) | 25.62 (13.96) | 35.84 (31.72) | 25.56 (13.18) |
| BT47 | SKBR | **6.094** (2.939) | 7.293 (1.989) | 6.304 (2.849) | 7.172 (3.929) | 17.50 (5.528) | 7.548 (3.947) | 5.810 (3.341) | SHSY | SKBR | 7.385 (2.325) | 6.836 (2.253) | **6.488** (2.665) | 6.599 (2.668) | 13.41 (2.069) | 9.155 (4.839) | 5.810 (3.341) |
| BT47 | SKOV | **12.50** (6.176) | 14.10 (2.297) | 13.76 (8.097) | 39.15 (30.45) | 29.60 (4.807) | 39.02 (37.53) | 26.63 (31.72) | SHSY | SKOV | 13.44 (2.777) | 15.05 (3.119) | **12.05** (2.775) | 14.16 (2.068) | 22.62 (3.493) | 24.89 (17.99) | 26.63 (31.72) |
| BV2 | A172 | 50.08 (17.13) | 11.50 (2.347) | 27.37 (10.35) | **11.14** (1.896) | 78.97 (44.53) | 27.00 (24.73) | 12.35 (6.537) | SKBR | A172 | 16.06 (13.07) | 21.55 (14.71) | **15.34** (10.41) | 17.69 (14.72) | 20.85 (13.35) | 15.46 (10.51) | 12.35 (6.537) |
| BV2 | BT47 | 68.42 (30.95) | 24.43 (6.746) | 37.08 (17.02) | 22.17 (9.288) | **21.13** (6.260) | 22.82 (10.08) | 17.91 (7.316) | SKBR | BT47 | 22.89 (10.42) | 27.59 (8.549) | 23.11 (10.40) | 23.04 (10.21) | 24.63 (10.14) | **22.53** (10.23) | 17.91 (7.316) |
| BV2 | Huh7 | 127.8 (43.61) | 31.36 (15.41) | 81.54 (32.62) | 28.39 (14.29) | 46.02 (14.46) | **22.67** (11.30) | 25.46 (14.63) | SKBR | BV2 | 7.020 (6.529) | **6.249** (3.964) | 6.865 (5.262) | 6.319 (4.830) | 13.32 (5.377) | 8.189 (7.388) | 5.574 (5.844) |
| BV2 | MCF7 | 48.64 (31.26) | **13.45** (6.483) | 23.57 (16.05) | 28.76 (20.36) | 19.65 (6.465) | 35.09 (29.97) | 25.56 (13.18) | SKBR | Huh7 | **26.76** (14.87) | 27.73 (14.14) | 28.04 (15.57) | 26.84 (14.54) | 39.22 (19.67) | 33.22 (15.45) | 25.46 (14.63) |
| BV2 | SHSY | 21.72 (6.459) | **8.618** (2.953) | 18.67 (7.034) | 10.04 (6.004) | 34.67 (16.68) | 21.13 (9.782) | 6.468 (2.137) | SKBR | MCF7 | 32.50 (23.24) | **31.75** (21.82) | 33.55 (25.33) | 34.99 (28.75) | 36.73 (28.83) | 34.04 (26.51) | 25.56 (13.18) |
| BV2 | SKBR | **6.812** (2.259) | 8.312 (2.615) | 7.866 (2.921) | 7.795 (2.126) | 17.58 (4.554) | 7.620 (3.672) | 5.810 (3.341) | SKBR | SHSY | 11.71 (3.322) | 15.20 (8.303) | 11.31 (5.614) | **9.024** (6.058) | 20.76 (8.719) | 17.47 (9.255) | 6.468 (2.137) |
| BV2 | SKOV | 60.52 (12.99) | **14.84** (2.518) | 42.78 (12.60) | 18.94 (8.271) | 78.57 (22.33) | 25.80 (3.681) | 26.63 (31.72) | SKBR | SKOV | 16.12 (12.45) | 19.01 (9.445) | **13.14** (3.904) | 20.50 (13.86) | 23.92 (11.11) | 16.25 (2.380) | 26.63 (31.72) |
| Huh7 | A172 | 16.63 (12.86) | 40.99 (32.48) | **16.45** (11.16) | 19.52 (12.89) | 41.13 (26.52) | 19.27 (15.51) | 12.35 (6.537) | SKOV | A172 | 21.69 (21.39) | 24.39 (25.55) | 26.70 (25.17) | **20.98** (19.34) | 51.44 (52.21) | 23.99 (23.51) | 12.35 (6.537) |
| Huh7 | BT47 | **20.24** (7.779) | 34.11 (25.71) | 20.40 (8.439) | 22.11 (9.146) | 33.34 (13.50) | 22.43 (10.15) | 17.91 (7.316) | SKOV | BT47 | 26.85 (10.92) | 50.51 (32.12) | 28.16 (10.38) | **24.67** (10.95) | 35.71 (22.80) | 25.01 (9.766) | 17.91 (7.316) |
| Huh7 | BV2 | **6.894** (5.257) | 170.8 (158.5) | 20.12 (19.60) | 12.31 (8.943) | 39.68 (32.19) | 23.47 (18.61) | 5.574 (5.844) | SKOV | BV2 | **10.63** (6.675) | 18.03 (18.01) | 13.87 (8.579) | 13.06 (10.43) | 27.06 (11.92) | 10.84 (8.609) | 5.574 (5.844) |
| Huh7 | MCF7 | 27.68 (20.09) | 55.21 (47.80) | 28.70 (20.36) | 32.91 (24.32) | **26.70** (7.547) | 27.83 (14.70) | 25.56 (13.18) | SKOV | Huh7 | 35.08 (16.07) | 56.17 (15.82) | 36.95 (16.55) | **34.26** (16.61) | 41.30 (15.91) | 37.68 (15.52) | 25.46 (14.63) |
| Huh7 | SHSY | **8.568** (2.024) | 34.95 (14.25) | 15.05 (3.746) | 15.18 (5.386) | 33.67 (16.04) | 20.20 (6.024) | 6.468 (2.137) | SKOV | MCF7 | 36.75 (32.08) | **22.38** (8.297) | 37.14 (32.02) | 35.24 (28.64) | 45.78 (39.79) | 37.03 (31.88) | 25.56 (13.18) |
| Huh7 | SKBR | **6.072** (2.679) | 11.44 (4.667) | 8.511 (4.240) | 9.392 (5.209) | 30.90 (15.21) | 9.400 (4.761) | 5.810 (3.341) | SKOV | SHSY | **10.01** (6.263) | 37.59 (15.58) | 13.20 (7.726) | 16.77 (9.587) | 44.74 (31.79) | 21.08 (13.11) | 6.468 (2.137) |
| Huh7 | SKOV | **14.26** (9.189) | 21.73 (6.484) | 27.15 (25.78) | 15.57 (4.373) | 38.60 (17.96) | 29.52 (32.24) | 26.63 (31.72) | SKOV | SKBR | 8.991 (4.256) | 13.87 (3.827) | 10.76 (5.797) | **8.551** (4.773) | 26.09 (14.16) | 10.46 (5.399) | 5.810 (3.341) |

Table S5: Comparison of segmentation performance for average surface distance (ASD)
Values in the table represent the mean ASD for the test data, and those in parentheses show the standard deviation. Those in bold type represent the model with the lowest ASD, except for the upper-bound model. Underlines represent the models with lower ASD than the upper-bound model. The lower-bound model is the model trained with only source data. The upper-bound model is the model trained with only target data. To save space, the names of the source and target cell lines have been abbreviated as follows: BT-474 to BT47, BV-2 to BV2, SH-SY5Y to SHSY, SK-BR-3 to SKBR, and SK-OV-3 to SKOV.

| Source | Target | Lower bound | Saito | Zou | Na | Wang | Ours | Upper bound | Source | Target | Lower bound | Saito | Zou | Na | Wang | Ours | Upper bound |
|---|---|---|---|---|---|---|---|---|---|---|---|---|---|---|---|---|---|
| A172 | BT47 | 5.491 (2.764) | 11.08 (2.811) | 5.829 (3.402) | **5.414** (3.014) | 8.742 (4.249) | 6.273 (3.191) | 3.907 (1.759) | MCF7 | A172 | 3.768 (3.243) | 3.297 (1.312) | 3.425 (2.692) | 4.112 (3.735) | 9.929 (4.576) | **3.230** (1.669) | 2.988 (1.774) |
| A172 | BV2 | 2.051 (1.120) | 2.506 (1.403) | 2.806 (1.171) | **1.911** (1.067) | 5.700 (1.357) | 11.44 (8.661) | 1.370 (0.967) | MCF7 | BT47 | **4.245** (1.991) | 5.081 (2.392) | 5.662 (3.064) | 4.835 (2.441) | 7.622 (2.503) | 5.282 (2.937) | 3.907 (1.759) |
| A172 | Huh7 | 6.362 (2.186) | 22.59 (7.554) | **5.309** (2.815) | 5.905 (2.934) | 9.994 (3.814) | 9.386 (3.699) | 4.691 (2.506) | MCF7 | BV2 | **1.909** (1.318) | 2.259 (1.259) | 2.173 (1.631) | 2.001 (1.206) | 5.386 (1.107) | 2.284 (1.023) | 1.370 (0.967) |
| A172 | MCF7 | **9.974** (10.66) | 10.54 (11.50) | 10.37 (11.46) | 10.37 (11.40) | 12.18 (10.54) | 10.52 (11.47) | 6.459 (4.175) | MCF7 | Huh7 | 5.836 (2.379) | 6.711 (2.506) | 5.919 (2.447) | **5.109** (2.334) | 11.40 (4.118) | 6.362 (3.161) | 4.691 (2.506) |
| A172 | SHSY | **2.400** (0.631) | 3.862 (1.801) | 2.506 (1.063) | 2.552 (1.051) | 8.685 (4.793) | 13.74 (11.18) | 1.508 (0.546) | MCF7 | SHSY | 2.450 (0.688) | 3.305 (1.513) | 2.541 (0.893) | **2.423** (1.011) | 9.326 (5.081) | 9.050 (5.703) | 1.508 (0.546) |
| A172 | SKBR | **1.529** (0.514) | 1.954 (0.981) | 1.672 (0.818) | 1.803 (0.891) | 5.784 (2.129) | 2.779 (1.237) | 1.261 (0.516) | MCF7 | SKBR | **1.336** (0.432) | 1.749 (0.599) | 1.476 (0.545) | 1.530 (0.643) | 5.985 (1.501) | 1.551 (0.583) | 1.261 (0.516) |
| A172 | SKOV | 4.664 (1.212) | 5.659 (4.100) | **4.454** (4.970) | 8.183 (8.648) | 12.54 (8.909) | 10.65 (15.07) | 6.357 (9.391) | MCF7 | SKOV | 4.443 (1.425) | **3.616** (1.169) | 3.802 (0.988) | 5.475 (3.380) | 8.444 (1.850) | 4.529 (1.139) | 6.357 (9.391) |
| BT47 | A172 | 3.466 (3.074) | 3.747 (1.012) | 3.282 (2.455) | 4.213 (3.540) | 6.825 (1.779) | **3.209** (2.008) | 2.988 (1.774) | SHSY | A172 | 4.073 (1.048) | 2.922 (0.710) | **2.524** (0.841) | 3.266 (1.868) | 7.276 (2.739) | 4.128 (3.968) | 2.988 (1.774) |
| BT47 | BV2 | **1.871** (0.927) | 4.066 (1.724) | 2.228 (0.935) | 2.307 (1.284) | 5.269 (1.308) | 3.208 (1.198) | 1.370 (0.967) | SHSY | BT47 | 8.625 (3.583) | 7.521 (2.098) | **3.697** (1.466) | 3.882 (1.647) | 7.543 (2.794) | 5.061 (2.704) | 3.907 (1.759) |
| BT47 | Huh7 | 5.233 (2.296) | 5.626 (2.139) | **4.925** (2.448) | 5.380 (2.700) | 8.583 (2.715) | 5.611 (2.967) | 4.691 (2.506) | SHSY | BV2 | **1.503** (0.752) | 1.786 (0.803) | 1.653 (0.881) | 1.809 (0.976) | 5.196 (1.645) | 1.961 (1.138) | 1.370 (0.967) |
| BT47 | MCF7 | **5.402** (2.977) | 5.790 (3.854) | 7.411 (5.422) | 8.194 (6.878) | 7.605 (3.280) | 9.782 (10.46) | 6.459 (4.175) | SHSY | Huh7 | 24.46 (10.68) | 6.665 (2.469) | **5.452** (2.080) | 5.710 (2.284) | 9.443 (3.022) | 5.813 (2.831) | 4.691 (2.506) |
| BT47 | SHSY | **2.071** (0.726) | 2.808 (0.682) | 2.788 (0.757) | 3.068 (0.860) | 6.982 (1.294) | 5.224 (3.312) | 1.508 (0.546) | SHSY | MCF7 | 5.999 (3.252) | 5.918 (3.386) | **5.253** (2.775) | 10.04 (10.95) | 8.537 (5.067) | 10.21 (11.27) | 6.459 (4.175) |
| BT47 | SKBR | **1.313** (0.466) | 1.745 (0.420) | 1.356 (0.450) | 1.489 (0.665) | 5.494 (0.727) | 1.549 (0.680) | 1.261 (0.516) | SHSY | SKBR | 1.577 (0.396) | 1.571 (0.422) | **1.390** (0.421) | 1.392 (0.433) | 4.694 (0.550) | 1.874 (0.924) | 1.261 (0.516) |
| BT47 | SKOV | **3.039** (1.657) | 3.762 (0.832) | 3.297 (2.196) | 11.79 (11.37) | 10.26 (1.283) | 10.21 (13.19) | 6.357 (9.391) | SHSY | SKOV | 3.296 (0.875) | 3.882 (0.927) | **2.937** (0.880) | 4.048 (0.800) | 7.845 (1.073) | 6.414 (5.436) | 6.357 (9.391) |
| BV2 | A172 | 16.03 (7.319) | 3.455 (0.876) | 7.086 (3.414) | **3.068** (0.747) | 30.46 (19.56) | 7.266 (8.269) | 2.988 (1.774) | SKBR | A172 | 3.807 (3.774) | 4.932 (3.487) | **3.568** (3.032) | 4.370 (4.714) | 6.609 (4.822) | 3.646 (3.083) | 2.988 (1.774) |
| BV2 | BT47 | 24.29 (13.75) | 6.071 (1.939) | 9.860 (5.368) | 5.411 (2.709) | 7.317 (1.868) | **5.334** (2.877) | 3.907 (1.759) | SKBR | BT47 | 5.232 (2.909) | 6.525 (2.922) | 5.327 (2.931) | 5.292 (2.847) | 7.739 (3.560) | **5.164** (2.834) | 3.907 (1.759) |
| BV2 | Huh7 | 49.16 (20.01) | 7.656 (2.424) | 26.61 (12.82) | **6.544** (2.520) | 14.96 (2.787) | 7.186 (2.067) | 4.691 (2.506) | SKBR | BV2 | 1.705 (1.100) | 1.630 (0.859) | 1.686 (0.948) | **1.595** (0.867) | 4.589 (1.454) | 1.948 (1.356) | 1.370 (0.967) |
| BV2 | MCF7 | 14.98 (13.78) | **3.750** (1.574) | 6.069 (4.707) | 7.975 (7.065) | 6.820 (1.536) | 10.11 (10.78) | 6.459 (4.175) | SKBR | Huh7 | **5.001** (2.764) | 6.640 (2.633) | 5.367 (2.785) | 5.234 (2.472) | 9.983 (4.390) | 6.204 (2.995) | 4.691 (2.506) |
| BV2 | SHSY | 6.158 (1.931) | **2.150** (0.584) | 4.861 (1.578) | 2.398 (0.777) | 10.85 (4.923) | 5.906 (3.120) | 1.508 (0.546) | SKBR | MCF7 | **8.977** (8.240) | 9.002 (8.183) | 9.505 (9.300) | 10.02 (10.76) | 11.74 (11.24) | 9.589 (9.256) | 6.459 (4.175) |
| BV2 | SKBR | **1.535** (0.400) | 1.988 (0.586) | 1.705 (0.476) | 1.851 (0.499) | 5.457 (1.077) | 1.577 (0.652) | 1.261 (0.516) | SKBR | SHSY | 2.810 (0.855) | 3.921 (2.227) | 2.618 (0.859) | **2.049** (0.842) | 6.993 (3.198) | 4.518 (2.593) | 1.508 (0.546) |
| BV2 | SKOV | 19.64 (4.445) | **4.067** (0.896) | 12.75 (4.743) | 4.881 (1.422) | 27.92 (7.249) | 9.317 (1.529) | 6.357 (9.391) | SKBR | SKOV | 3.717 (3.208) | 4.780 (2.504) | **3.241** (1.163) | 5.223 (4.226) | 7.918 (3.732) | 4.893 (0.942) | 6.357 (9.391) |
| Huh7 | A172 | 3.920 (3.633) | 9.859 (8.294) | **3.791** (2.980) | 4.704 (3.918) | 13.47 (10.64) | 4.646 (4.840) | 2.988 (1.774) | SKOV | A172 | 5.346 (6.761) | 6.292 (8.533) | 6.843 (8.228) | **5.212** (6.269) | 17.94 (22.40) | 6.029 (7.609) | 2.988 (1.774) |
| Huh7 | BT47 | **4.492** (2.032) | 8.758 (9.991) | 4.538 (2.165) | 5.049 (2.551) | 10.47 (4.283) | 5.173 (2.818) | 3.907 (1.759) | SKOV | BT47 | 6.709 (3.720) | 16.49 (14.56) | 7.372 (3.805) | 5.997 (3.411) | 12.28 (9.515) | **5.961** (3.049) | 3.907 (1.759) |
| Huh7 | BV2 | **1.715** (0.936) | 72.25 (88.42) | 4.426 (5.360) | 3.181 (2.370) | 10.06 (6.923) | 4.125 (2.370) | 1.370 (0.967) | SKOV | BV2 | **2.672** (1.405) | 4.630 (4.763) | 3.189 (1.452) | 3.344 (2.423) | 7.907 (2.611) | 2.764 (2.013) | 1.370 (0.967) |
| Huh7 | MCF7 | **6.877** (5.861) | 15.98 (16.64) | 7.342 (6.351) | 9.302 (8.956) | 8.095 (2.028) | 7.477 (4.932) | 6.459 (4.175) | SKOV | Huh7 | 6.851 (3.496) | 12.64 (5.185) | 7.039 (3.416) | **6.365** (3.276) | 10.68 (4.165) | 7.083 (3.198) | 4.691 (2.506) |
| Huh7 | SHSY | **2.020** (0.549) | 9.247 (3.668) | 3.626 (0.991) | 3.886 (1.529) | 10.20 (3.747) | 5.716 (1.987) | 1.508 (0.546) | SKOV | MCF7 | 10.61 (11.50) | **6.487** (2.352) | 10.86 (11.57) | 10.18 (10.60) | 15.19 (14.92) | 10.73 (11.47) | 6.459 (4.175) |
| Huh7 | SKBR | **1.331** (0.442) | 2.479 (0.749) | 1.752 (0.754) | 1.900 (0.997) | 8.637 (3.013) | 1.945 (0.938) | 1.261 (0.516) | SKOV | SHSY | **2.245** (0.942) | 10.23 (5.113) | 3.152 (1.624) | 3.977 (2.702) | 15.43 (12.65) | 5.424 (3.914) | 1.508 (0.546) |
| Huh7 | SKOV | **3.404** (2.264) | 5.594 (1.896) | 6.522 (7.891) | 3.936 (1.324) | 12.56 (5.918) | 7.295 (9.963) | 6.357 (9.391) | SKOV | SKBR | 1.894 (0.785) | 3.985 (0.435) | 2.303 (1.152) | **1.752** (0.869) | 8.220 (4.332) | 2.255 (1.108) | 1.261 (0.516) |
